# Agricultural intensification favours an introduced bumble bee over its native congener through differences in foraging range, habitat association, and lineage continuity

**DOI:** 10.64898/2026.05.07.723627

**Authors:** Jenna B. Melanson, Tyler T. Kelly, Natalia Clermont, Jonathan B. Uhuad Koch, Claire Kremen

## Abstract

Agricultural intensification can enhance the expansion of introduced species which are highly adapted to human-modified landscapes, but the mechanisms by which this occurs are often unclear. Here we investigate the spatial ecology of a rapidly expanding introduced bumble bee (*Bombus impatiens*) and a native congener (*B. mixtus*) in agricultural landscapes of southwestern British Columbia, Canada. We used microsatellite genotyping and spatially explicit capture-recapture models to compare the foraging distance of the two species, and fitted hierarchical models to compare their abundance, behaviour (nest searching vs foraging), and lineage survival as a function of landscape composition and configuration. We found that *B. impatiens* had a broader foraging range than *B. mixtus*. *B. impatiens*’ colony/worker abundance were positively associated with the surrounding area of residential gardens and field edges, but decreased relative to *B. mixtus* abundance in response to increasing seminatural area. In contrast, *B. mixtus* colony abundance decreased in landscapes with a greater area of intensively managed berry crops. We observed fewer *B. impatiens* queens per survey in landscapes with more low-disturbance landcover, and hypothesize space use of this species could be shaped by concentration on potential nesting habitat. Consistent with this observation, nest searching behaviour was more common for *B. impatiens* queens, while *B. mixtus* queens varied in their use of certain habitat types for nest searching and were primarily observed foraging, suggesting these two species derive different value from agricultural landscapes during colony establishment. Finally, we found that the rate of lineage re-capture between 2022 colonies and 2023 spring queens was nearly 10-fold higher for *B. impatiens* than for *B. mixtus*, indicating a greater capacity of the introduced species to complete its life cycle in agro-natural landscape mosaics. Our results suggest that differences in spatial ecology may contribute to the differential success of these two species in human-modified landscapes, and provide insight into the mechanisms by which land-use change shapes community composition.

**Graphical abstract:** Coloured diagrams of *B. mixtus* and *B. impatiens* are credited to Elaine Evans and the Xerces Society, with permission.

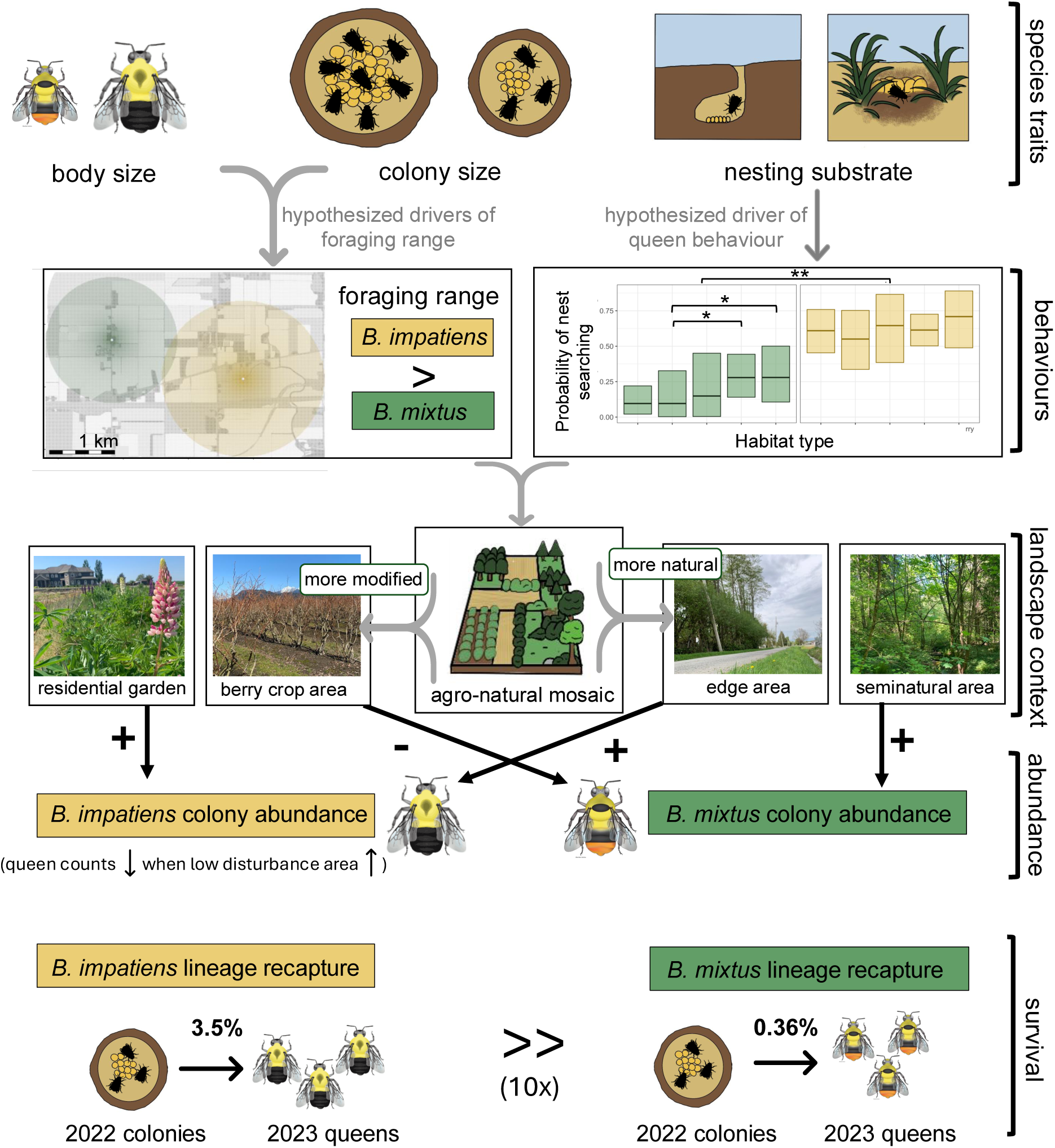

## 1 Introduction

Agricultural intensification shapes ecological communities through a variety of mechanisms, including habitat loss (Kremen et al., 2004), reduced landscape complexity (Martin et al., 2020; Sirami et al., 2019), exposure to pollutants (Geiger et al., 2010; Wan et al., 2025), and species introductions (Liu et al., 2023; Mooney & Hobbs, 2000). In agricultural landscapes, bee *α*- and *γ*-diversity are reduced by 10-20% globally in comparison to natural habitats (Tsang et al., 2025), a concerning pattern given the valuable contributions of wild bees to crop pollination (Dainese et al., 2019; Garibaldi et al., 2013). But the effects of agricultural intensification are not felt equally across taxa. Anthropogenic stressors can produce divergent population trends, with species sorting into so-called “winners” and “losers” depending on their tolerance for novel conditions (Dornelas et al., 2019; McKinney & Lockwood, 1999); when tolerance is related to species’ traits, this process can lead to the taxonomic or functional homogenization of pollinator communities (Fisogni et al., 2025). Identifying differences in the spatial or behavioural ecology of closely related winners and losers in a shared landscape context can provide insight into both the characteristics which lead to success in human-modified landscapes, as well as the management practices which mitigate or exacerbate species’ differential responses.

Bumble bees (*Bombus*) are critically important temperate pollinators which are integral to wild plant-pollinator networks (Memmott et al., 2004) and highly effective at pollinating many crops (Goulson, 2010). While there is growing evidence of declines for some bumble bee species (Cameron & Sadd, 2020; Kerr et al., 2015), other species are stable or expanding within and/or beyond their native ranges (Averill et al., 2021; Jackson et al., 2022). Although multiple global change drivers have been implicated in bumble bee declines, identifying when and how agricultural intensification shapes *Bombus* community composition—and which traits enable persistence in modified landscapes—could improve predictions of future communities, forecasts of pollination service availability, and the design of management practices to support vulnerable species.

Behavioural traits related to bumble bee spatial ecology are likely to be particularly important for understanding species’ responses to agricultural intensification. For example, a larger foraging range may allow species to locate and use highly abundant but temporally transient food sources in agricultural landscapes (Walther-Hellwig & Frankl, 2000), where removal of native vegetation and spatial/temporal concentration of floral resource availability in mass-flowering crops can lead to a more patchy distribution of resources than in natural landscapes. Previous work suggests that differences in foraging range permit coexistence amongst bumblebee species with similar dietary niches (Westphal et al., 2006), but mechanistic models investigating the trade-offs between foraging efficiency and travel costs indicate that such coexistence may only be possible when landscapes are compositionally and configurationally complex (Bolin et al., 2018). Landscape simplification may therefore be expected to tip the coexistence balance in favour of species with larger foraging ranges.

Another key aspect of bumblebee spatial ecology is the selection of nesting sites, which must meet species-specific microhabitat requirements, remain undisturbed throughout the colony cycle, and occur close enough to floral resources to support colony establishment, growth, and reproduction (Goulson, 2010). Species with less restrictive nesting requirements may be able establish nests across landcover types experiencing intermediate levels of disturbance, and may therefore exhibit different spatial patterns than those with more specialized nesting niches or those that nest in easily disturbed microhabitats.

Foraging range and nesting habitat suitability are relatively challenging traits/behaviours to measure in bumble bees, often captured through proxy measures such as queen nest searching surveys or spatial modelling of genetic capture-recapture data. While it would be ideal to study such traits at a community level, it is more feasible to make targeted comparisons between species which we expect to vary in their spatial ecology. Here, we draw comparisons between a highly abundant introduced/invasive species *Bombus impatiens* (the Common Eastern Bumble Bee) and its native congener, *B. mixtus* (the Fuzzy-Horned Bumble Bee) in agricultural landscapes of southwestern British Columbia (Canada). Both species are considered common in this region, but *B. mixtus* exhibits a stable population size, while *B. impatiens* is expanding rapidly throughout the Pacific Northwest after its escape from greenhouse facilities in the early 2000s (Looney et al., 2019).

By comparing *B. mixtus* and *B. impatiens* we investigate how bumble bee space and habitat use are related to spatial patterns of abundance and lineage turnover in agroecosystems. Where possible, we make use of community-level data to support our conclusions. Specifically we ask:

1. How does foraging distance vary between a highly successful introduced and a relatively common native species?
2. How do the colony, worker, and queen abundances of the two species relate to availability of foraging and nesting habitat?
3. How frequently do queens of the two species utilize different habitat types for nest-searching versus foraging? Are these trends consistent throughout the *Bombus* community?
4. How do agricultural landscapes support the reproduction/lineage continuity of *B. mixtus* and *B. impatiens*?

Beyond contrasting the space use and land-use responses of these two species, our work provides insight into the ecology of a potentially invasive species in its novel range, including habitat associations that may inform efforts to predict its future spread.

## 2 Methods

### 2.1 Study system

The study took place in the Lower Fraser Valley of southwestern British Columbia, Canada, in an agricultural system dominated by hay, mixed vegetable, and perennial berry production (Fig. A1). Field surveys were conducted across six replicate landscapes, each encompassing roughly 3km^2^ of farmland interspersed with rural/suburban residence (Fig. A2). In each landscape, we established 30 semi-evenly dispersed sampling transects (50 meters x 2 meters); the size of landscapes and spacing of transects were designed to facilitate recaptures of colonymates at multiple spatial locations, while covering a large enough area to observe longer-distance foraging events (up to 2.5 km apart within a given landscape). Hereafter we refer to these spatial divisions as “landscapes” and “transects.”

We conducted 10 sampling rounds for each landscape from May-August 2022 and 17 sampling rounds from March-August 2023; while transect turnover occurred within and between years due to changes in land access, the number of surveys per landscape per sampling round varied little (28.1 ± 1.9 (mean ± SD) in 2022, 30.2 ± 0.9 in 2023). Throughout the analyses we incorporate offset variables when relevant to account for differences in survey effort per transect. A total of 289 transects (28,900*m*^2^) were visited during 395 total person hours of sampling over the course of the study.

### 2.2 Study species

*Bombus impatiens* is native to eastern North America and has expanded rapidly throughout southwestern BC in recent decades, where it overlaps with native *B. mixtus*. Both species follow an annual life cycle: queens emerge from hibernation in spring, forage and search for nesting sites (March–May), and establish colonies from which daughter-workers take over foraging duties through summer (May–August). In late summer, colonies produce reproductive individuals (males and gynes) which leave the nest and mate; colonies senesce, and fertilized queens go into hibernation to begin the cycle anew in the following year (Williams et al., 2014). We targeted both species across the worker foraging period in 2022 and 2023, and during the queen emergence and colony establishment phase in 2023.

### 2.3 *Bombus* collections and floral surveys

*Bombus* surveys at each transect entailed 5 minutes of active search time (totaling 140 hours in 2022 and 255 hours in 2023), during which the stopwatch was paused whenever a foraging bumble bee was sighted. Workers were captured by netting, placed into sterile 15 mL tubes, and immediately placed on ice before transfer to a -80°C freezer at the end of the day. We collected tarsal clips from queens and released them at the location of capture. We recorded each queen as either foraging (visiting flowers) or nest searching (flying low to the ground in a zig-zag pattern, or crawling on the ground to investigate nest sites). In 2022 and 2023 we collected workers of both species; in 2023, queen observations were made for the full *Bombus* community, but only queens from the focal species (*B. mixtus* and *B. impatiens*) were sampled for genotyping. We sampled from May-August in 2022, and March-August in 2023, to capture relevant stages of the *Bombus* life cycle. The long duration of sampling in 2023 (March-August) provided the opportunity to observe both spring queens (originating in 2022) and summer gynes (originating in 2023); to establish a cutoff between these groups, we visually inspected histograms of queen counts per day and chose thresholds of 15 June and 26 June for *B. mixtus* and *B. impatiens*, respectively (Fig. A3). Queens were assigned as spring queens if they were observed before these dates, and as summer gynes if they were observed after. Surveys were conducted on days when the temperature was above 12^◦^ C (10^◦^ C for queen surveys) and wind speeds below 2.5 m/s.

We identified all flowering plants within each 50 m x 2 m transect to species or genus using the iNaturalist Seek mobile application (Hart et al., 2023). Inflorescence abundance was visually estimated in the field for each species along each transect. Estimates were assigned to one of five log-scale abundance categories (0-4: 0 = 1-10, 1 = 11-100, 2 = 101-1,000, 3 = 1,001-10,000, 4 ≥ 10,000) (Cariveau et al., 2024). Floral surveys were filtered to exclude species never visited by bumble bees, removing noise from highly abundant but small-flowered herbs unattractive to *Bombus* foragers.

### 2.4 Landscape characterization

Land cover maps were developed for each study landscape based on manual classification of Google Earth satellite imagery. Raster files were downloaded into QGIS (QGIS Development Team, 2024), and a polygon-based vector layer denoting land classes was created via visual inspection by a single observer (TTK) familiar with the study system. Classification was at 2-meter resolution and utilized 17 categories: annual row crops, blueberry, cranberry, other perennial crops, crop polycultures, hay meadows, pasture, fallow, grassy field margins, hedgerows (tree-dominated), hedgerows (blackberry-dominated); seminatural grassland, forest, wetlands; residential gardens, impervious surfaces, and water. These land cover categories were chosen based on their hypothesized provisioning of floral resources and different disturbance regimes (e.g., tilling/flooding/mowing) which could impact survival of subterranean or surface-level bumble bee nests. The vector layer was converted into a raster file and subsequent analyses were performed in R (R Core Team, 2021).

To quantify landscape configuration, we used the interspersion and juxtaposition index (IJI) as computed in the *landscapemetrics* package (Villalba et al., 2024); this metric measures how evenly patches of different classes are interspersed with one another based on their shared edge length, and is presented as a percent of the maximal ad-jacency for a given set of classes (McGarigal & Marks, 1995). We chose IJI because it was identified by Kennedy et al. (2013) as a configurational metric most orthogonal to measures of landscape composition. For queen abundance models, landscape composition was also quantified using the *landscapemetrics* package. For colony abundance models, we computed the distance-weighted area of each type of foraging habitat; see below for methodology.

### 2.5 PCR Amplification and Fragment Analysis

DNA was extracted from the mid-leg basitarsus and tarsus of each worker (distal tarsus only for queens) using the HOTSHOT protocol (Truett et al., 2000). Specimens were microsatellite genotyped at loci BT10, BTERN01, BL13, BL15, B126, BTMS0057, BTMS0059, BTMS0062, BTMS0083 (both species); BTMS0066, BTMS0086, BTMS0104, BTMS0126, BTMS0136 (*B. mixtus* only); BT28, BT30, B10, B96, B124, BTMS0073, and BTMS0081 (*B. impatiens* only) (Estoup et al., 1995, 1996; Reber Funk et al., 2006; Stolle et al., 2009). For details of the genotyping protocol, see Appendix Section A2.1. Microsatellite peaks were assigned manually using the software Geneious Prime 2024.0.5 (https://www.geneious.com). Scoring error rates were assessed by re-genotyping a panel of 96 individuals per species, and a locus with observed error rate ≥3% was discarded from further analyses (BL15 for both species).

### 2.6 Colony Assignments

We performed locus quality filtering based on deviations in *F_is_*estimates across loci, and linkage disequilibrium between locus pairs (Appendix Section A2). After removal of deviating loci (BTERN01, BTMS0104, and BTMS0059 for *B. mixtus*, BTMS0073, BT28, BL13 for *B. impatiens*) we performed colony assignments for individuals which were successfully genotyped at ≥ 8 microsatellite loci. Samples with fewer than 8 scored loci were rare and randomly distributed across transects/years/sampling rounds.

We performed colony assignments using the pedigree reconstruction software COLONY 2.0.6.5 (Jones & Wang, 2010). To identify the optimal software settings for our dataset, we performed a simulation study using allele frequencies derived from our true data. We found that improper choice of software parameterization can lead to unacceptably high rates of assignment of unrelated individuals to colony groups, in some instances resulting in a false positive rate of up to 60%; however our chosen software setting (described below and discussed in Appendix Section A2.3) achieved false positive and false negative rates of ≤ 5% for simulated data.

We assigned colonies separately for each species-year combination, with exclusion tables to prevent assignment of siblingships between individuals originating from different landscapes. Each run of the software assumed male and female monogamy and no inbreeding, and used no prior on siblingship size and no siblingship size scaling. Runs were performed using the Linux command line version of the software, with run length set to “long”, likelihood set to “full-likelihood” and precision set to “high precision.” Sibling pairs were maintained with *P_f ullsibdyad_* = 0.995. Because we used full sibling dyad probabilities rather than sibling cluster probabilities to perform assignments, we resolved non-circular families as described in A2.3.3. We included 2023 spring queens in the 2022 software run, as these queens were putative siblings of 2022 workers. We also identified mother-daughter relationships between 2023 spring queens and 2023 workers/gynes in a separate software run (see Fig. A4 for pedigree diagram).

### 2.7 Statistical Models

All analyses were carried out using R version 4.3.3 (R Core Team, 2021). Bayesian model inference was conducted in Stan (Carpenter et al., 2017) or *brms* (Bürkner, 2017); we ran foraging distance models on 4 chains of 5000 iterations each, where the first 1000 iterations of each chain were discarded as warmup; all other models were run for 6000 iterations (1000 warmup, 5000 sampling). We assessed the quality of inference by monitoring *R*^ convergence diagnostics and effective sample size. For models of abundance, behaviour, and lineage survival, we used posterior predictive checks to confirm that fitted models generated predictions which closely matched our observed data, and chose error distributions which led to the closest alignment between real and model-predicted observations. For all models we checked pairwise Pearson product-moment correlation between predictors before model-fitting and ensured that all correlation coefficient magnitudes were less than 0.5. Finally, we tested model residuals using Moran’s I at two neighborhood scales (500m and 1000m) and found no evidence of spatial autocorrelation (all p-values > 0.05) (Table A8).

In the text and figures we present marginal predicted probabilities/counts, obtained by averaging posterior expected predictions over the observed distribution of covariates. To describe the effect size of predictors of interest, we computed the average marginal effect (AME) across observed covariate values. We prefer the AME to model summary coefficients because the AME is on the outcome scale (rather than linear predictor scale) and is therefore more biologically interpretable. We calculate AMEs by applying a one standard deviation increase in the predictor for each observation in the dataset and computing the average change in the predicted outcome across observations; averaging is necessary because changes to linear predictors on the log/logit scale result in nonlinear changes on the outcome (count/probability) scale. While total survey effort varied for colony, worker, and queen abundance models, we standardize AMEs by reporting the average marginal change in the response variable that would be expected in 60 minutes of survey time. Main figures show marginalized predictions across transects varying in survey effort (colony, worker abundance models) and per-survey predictions (queen abundance models). We consider an association to have “strong support” when 95% credible intervals for the AME exclude zero, and “mod-erate support” when 90% credible intervals excluded zero. Conditional effects on the linear predictor scale are presented in Tables A1-A7.

#### 2.7.1 Foraging Distance

To compare foraging distance of the two species, we fitted spatially explicit genetic capture-recapture models following Pope and Jha (2017). These models parametrize the distance decay rate of foraging activity, conditioned on the observed frequency of workers from each colony (*i* ∈ *C*) at each transect (*k* ∈ *K*). We sum observations across sampling rounds and consider visitation events only within the landscape where a colony was observed (e.g., colony *i* can be observed only at transects *k* ∈ *K_l_*, where *K_l_* is the subset of transects occurring in the landscape where we observe *i*). The likelihood for a simple model where visitation intensity decays exponentially with distance is:

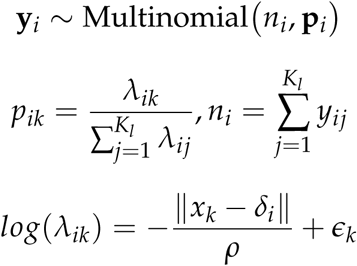

where *y_ij_* is the number of individuals from colony *i* captured at transect *k*, *δ_i_* is a nuisance parameter governing the latent location of colony *i*, *x_k_* − *δ_i_* is the Euclidean distance between transect *k* and colony *i*, *ρ* is the length scale of foraging activity, and *ɛ_k_* are transect-year-species intercepts drawn from a common distribution. We incur a sampling offset by adding the log-transformed sampling effort per transect on the linear predictor scale for each *λ_ik_*.

We fitted the model using a dataset including both *B. mixtus* and *B. impatiens* colonies, where *ρ* (the foraging length scale) was estimated independently for each species. We make comparisons of posterior estimates of *ρ* (and resulting visitation decay rates) for each species. To assess different assumptions about reasonable spatial distributions for colony locations *δ*, we apply bivariate Gaussian priors centered at the centroid of each replicate landscape. The results presented in the main body represent a scenario where 99% of observed colonies are located within 3km of the landscape centroid (i.e., within approx. 2.15 km of the edges of the transect grid). In Appendix Section A3, we perform sensitivity analyses on this prior, assuming 99% of colonies fall within 1.5km or 4.5km of the landscape centroid; in the same section, we further discuss model implementation and prior specification.

To facilitate comparison with previous studies (e.g., Jha and Kremen (2013), Mola et al. (2020), and Redhead et al. (2016)), we report summaries of the mean colony-specific foraging distance (i.e., the mean distance between each worker and its colony-specific centroid). These values do not take into account transect configuration or attractiveness. To account for sampling effort at each distance, we take a second, pairwise approach. We bin the minimum pairwise foraging distance (half the distance between sibling pairs) into 100 meter increments, and normalize the frequency of each bin by dividing by the number of transect-survey pairwise distances which fall within that bin (see Fig. 1A).

**Figure 1:**
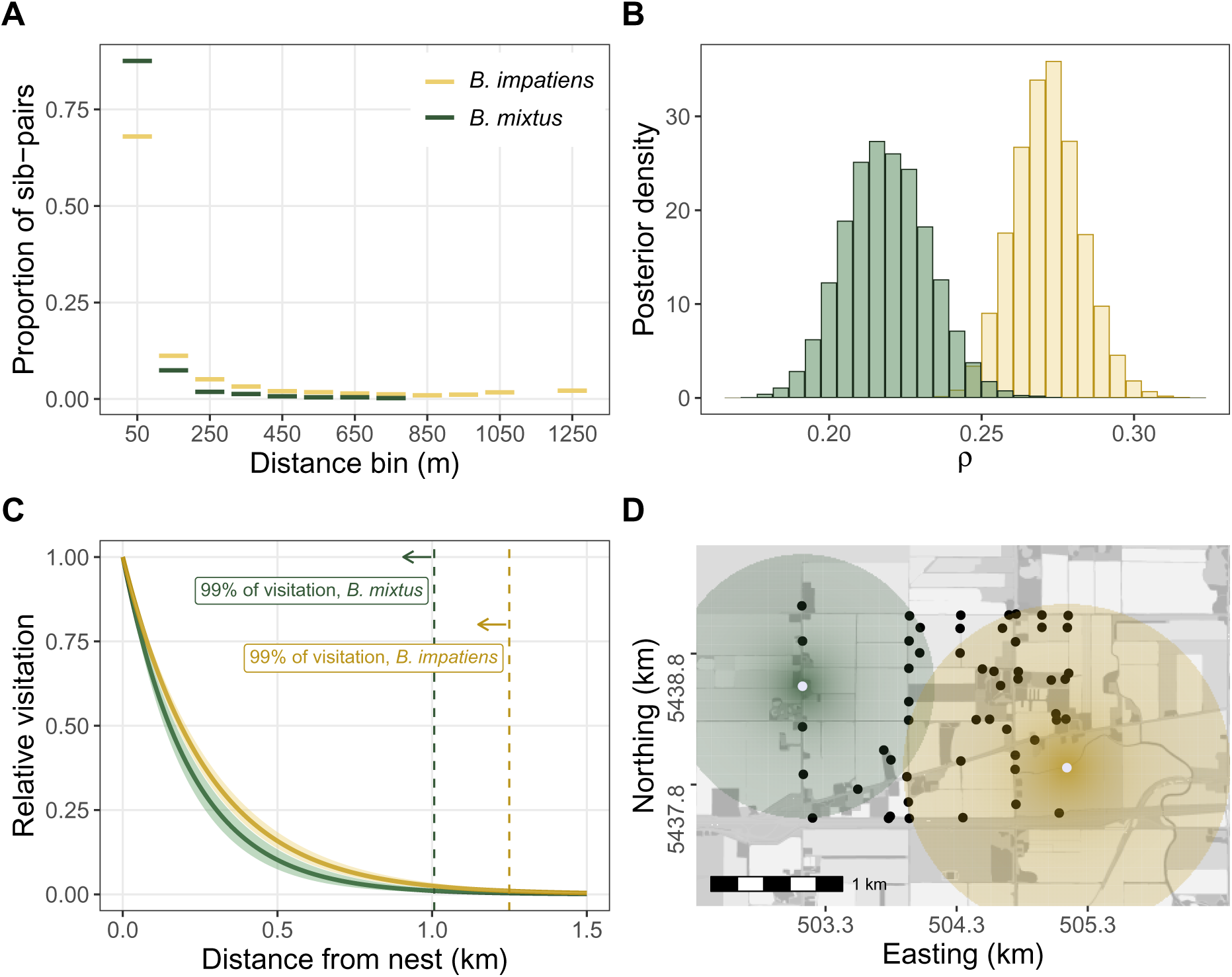
A) Observed pairwise minimum foraging distance for *B. mixtus* (green) and *B. impatiens* (yellow), normalized by the number of transect-survey pairs which could result in a foraging distance in each 100m bin. B) Posterior distributions of the foraging scale parameter (*ρ*) for each species, estimated from spatially explicit capture-recapture models. C) Model-predicted decay of visitation with increasing distance from the nest; horizontal dashed lines indicate the radius within which 99% of activity occurs. D) Decay kernels overlayed on one of our replicate landscapes. White points indicate theoretical colony locations (to center illustrative foraging kernels—not based on real data). Black points indicate true transect locations.

#### 2.7.2 Colony/worker abundance

We modelled the abundance of colonies and workers observed at each transect as a function of habitat composition and configuration at the scale of foraging for each species. We hypothesized that five habitat types had the greatest potential to provision colonies with floral resources, and would therefore be associated with colony or worker counts: seminatural area (forest, wetlands, and grasslands), field edge area (hedgerows and grassy margins), fallow area, perennial berry crops (blueberry and cranberry), and residential gardens.

For each transect we computed the distance-weighted area of each habitat type using the species-specific foraging scale (*ρ*) derived from models of foraging distance.

For each habitat type, the full landcover raster was converted to a binary raster (1 = focal habitat type, 0 = other); we then applied a weight to each pixel based on its Euclidean distance from the transect. To compute weights, we used posterior mean values of *ρ* for each species and the functional form of visitation decay described above. We summed the distance-weighted quantity of habitat within 4.6*ρ* of each transect (approx. 1000m for *B. mixtus* and 1200m for *B. impatiens*); any colony located beyond this distance would contribute an expected visitation intensity at the transect which was <1% of the visitation intensity at the colony location. We computed the interspersion and juxtaposition index (IJI) within the same buffer area (not distance-weighted).

We fitted models with colony/worker abundance (summed for each species across surveys within each year) as outcomes. As predictors we used the distance-weighted quantity of the five floral-provisioning habitat types described above, IJI, floral abundance (averaged across surveys), and year. Before fitting models, we centred and scaled each predictor independently for each species, to account for the different ranges of each predictor for each species (i.e., the number of distance weighted hectares of each landcover type is higher on average for a species with a larger foraging range). The effect of each predictor was estimated independently for each species. We included hierarchical intercepts for transect and landscape to account for correlation of multiple observations within each spatial grouping (often referred to as “random ef-fects”). We fitted models in *brms* using a negative binomial error distribution. We computed the average marginal effect (AME) of each predictor on each species, and quantified interaction contrasts by taking the difference of AME posteriors between species.

#### 2.7.3 Queen abundance

We modelled the abundance of nest-searching and foraging spring queens as a function of local- and landscape-scale nesting habitat, landscape configuration, floral abundance, and Julian date. We computed putative nesting habitat proportion at two scales (50m and 1000m) by combining landcover classes which we hypothesized could provide undisturbed subterranean and/or surface-level nest sites for bumble bees: hedgerows/field margins, blueberry and other perennials (excluding cranberry), fallow fields, forest, grassland, and residential gardens. Because we were not certain that this habitat was truly suitable for nesting for both species, we refer to it hereafter as “low disturbance habitat,” in reference to the lack of frequent tilling or flooding during the management of these habitat types. We modelled queen abundance for each species using survey-level data (i.e., a single survey on a single transect); because our dataset included up to 10 observations per transect prior to the cutoff dates, we included hierarchical intercepts for both transect and landscape. Based on posterior predictive checks we ascertained that the data were zero-inflated. We therefore used a zero-inflated Poisson error distribution with the zero-inflation term fitted as intercept-only.

#### 2.7.4 Queen behaviour

We next modelled the behaviour of *B. mixtus* and *B. impatiens* queens using a Bernoulli error distribution (1 = nest searching, 0 = foraging). We hypothesized that species identity, local habitat type, and floral abundance could play a role in determining queen behaviour. We classified each transect into one of four categories: field margin, roadside, blueberry field, or seminatural. We computed floral abundance during the survey in which each queen was observed. We allowed for independent effects of each habitat type and floral abundance on behaviour of each species. We used hierarchical intercepts for transect and landscape which were shared across species.

In support of our findings in *B. mixtus* and *B. impatiens*, we performed an additional analysis on the full *Bombus* community. We modelled the behaviour of queens of all observed *Bombus* species using the same model as above, but with complete pooling on habitat type and floral abundance predictors (i.e., not estimated separately per species) (Fig. A6, Table A6).

#### 2.7.5 Lineage re-capture

Spring queens are produced by the previous year’s colonies, making them siblings of the previous year’s workers. Here—as in our analysis of worker siblingships—we assume singly-mated queens and identify full siblingships only, although we acknowledge that this assumption can sometimes be violated for North American *Pyrobombus* (Payne et al., 2003). We assessed the rate of lineage re-capture between 2022 colonies and 2023 queens based on siblingship assignments between these groups; we used a Bernoulli distribution to model recapture of each colony (1 = colony recaptured, 0 = colony not recaptured). Because the number of lineage recaptures was low, we did not model recapture rate as a function of landscape covariates, but instead compared the recapture rates of *B. mixtus* and *B. impatiens* by using species as a predictor.

## 3 Results

We observed a total of 1,919 *B. mixtus* workers (2022: 762, 2023: 1,157) of which 1,883 (2022: 755, 2023: 1,128) were successfully genotyped and assigned to 1,677 colonies (2022: 673, 2023: 1,004). We observed a total of 3,361 *B. impatiens* workers (2022: 1,251, 2023: 2,110) of which 3,312 (2022: 1,213, 2023: 2,099) were successfully genotyped and assigned to 2,632 colonies (2022: 1,023, 2023: 1,609). The average number of individuals observed per colony was 1.12 for *B. mixtus* (172 colonies sampled with ≥ 2 siblings) and 1.26 for *B. impatiens* (435 colonies sampled with ≥ 2 siblings).

In 2023 we observed 508 queens (*B. mixtus*: 132, *B. impatiens*: 209, other species: 167 (Fig. A3)), of which 99 *B. mixtus* and 149 *B. impatiens* were captured and successfully genotyped.

### 3.1 Foraging distance

Foraging distance models supported a greater foraging range for *B. impatiens* than for *B. mixtus* (*ρ_impatiens_*− *ρ_mixtus_* = 53m, 95% CrI: [18, 86]) (Fig. 1). The approximate magnitude of this difference was robust to prior sensitivity analyses, although for more extreme prior specifications (e.g., 99% of colonies located within 1.5km or 4.5km of the landscape centroid) credible intervals slightly overlapped zero (Fig. A21). Our main model predicted that 99% of foraging activity occurs within 1007m (95% CrI: [885, 1138]) of the nest for *B. mixtus*, and within 1249m (95% CrI: [1150, 1355]) for *B. impatiens*. Our sensitivity analyses showed that estimates of absolute foraging distance are sensitive to prior specification (Fig. A21), meaning that these values should be interpreted with caution. We chose to use these estimates of foraging scale to inform distance-weighting of landscape metrics because they constitute our best approximation of foraging scale for these two species.

The average colony-specific foraging distance based on the siblingship centroid approach was 91.98 ± 11.98*m* (mean ± SE) for *B. mixtus* and 262.45 ± 12.35*m* for *B. impatiens*. The maximum observed distance between siblings was 1.48 km for *B. mixtus* and 2.45 km for *B. impatiens*.

### 3.2 Colony / worker abundance

We observed a strong positive association between floral abundance and colony abundance for both *B. mixtus* and *B. impatiens* (Fig 2A). A one standard deviation increase (i.e., 1.15 unit increase) in log-scaled floral abundance was accompanied by an increase of 2.72 (95% CrI: [1.52, 4.20]) *B. mixtus* colonies and 1.87 (95% CrI: [0.51, 3.48]) *B. impatiens* colonies per hour of survey time. There was no effect of landscape configuration on colony abundance of either species (Fig. 2B). We observed a negative association between the distance-weighted area of berry crops (blueberry/cranberry) and *B. mixtus* colony abundance; a one standard deviation increase in berry crop area was associated with a reduction of 1.44 (95% CrI: [0.73, 2.13]) *B. mixtus* colonies per hour. *B. impatiens* colony abundance also tended to decline with increasing berry crop area, but model support for the association was weak (Fig. 2C). In contrast, we found that a one standard deviation increase in residential garden area was associated with a gain of 1.18 (95% CrI: [−0.03, 2.57]) *B. impatiens* colonies per hour of survey time, although the support for this relationship was only moderate. There was no significant change for *B. mixtus* (Fig. 2D). There was moderate support for a positive effect of seminatural area on *B. mixtus* colony abundance (AME: 0.61, 95% CrI: [−0.08, 1.42]) and a negative but more weakly supported trend for *B. impatiens* (AME: -0.78, 95% CrI: [−1.67, 0.19]). There was strong evidence for a difference in the average marginal effects between species (*AME_mix_*− *AME_imp_* = 1.39, 95% CrI: [0.48, 2.29]) (Fig. 2E), supporting a shift in relative abundance of the two species in landscapes with more seminatural area. There was moderate support for an increase in *B. impatiens* colony abundance with increasing field margin area (AME: 1.10 (95% CrI: -0.18-2.54)); the trend for *B. mixtus* colony abundance was also positive, but posterior support was weak (Fig. 2F). There was little evidence of a trend for either species with respect to fallow area (Fig. 2G).

**Figure 2:**
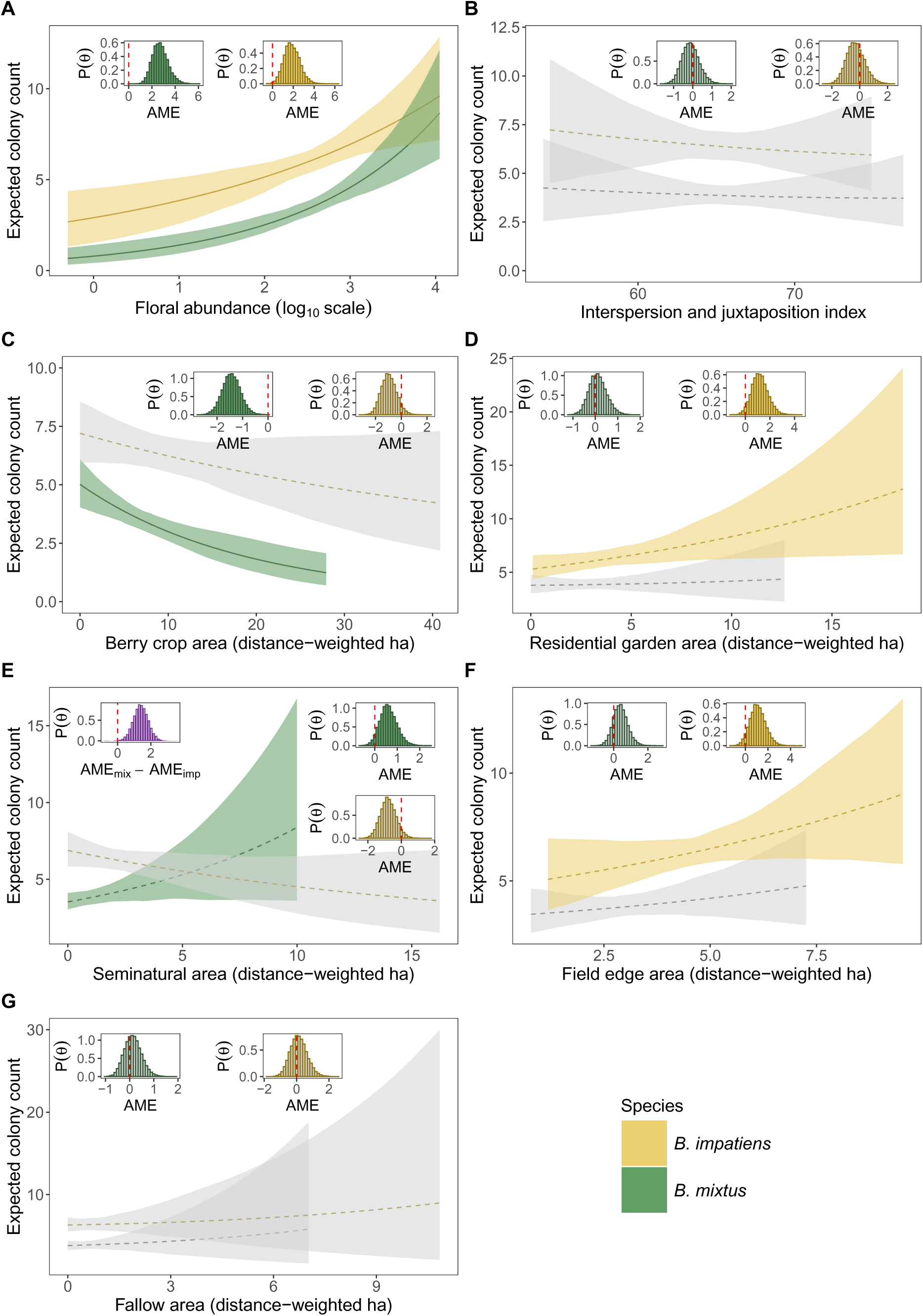
Colony abundance of *B. mixtus* (green) and *B. impatiens* (yellow) in relation to floral abundance (A), landscape interspersion (B), berry crop area (C), residential garden area (D), seminatural area (E), field edge area (F) and fallow arera (G). Each figure shows predicted colony counts across the range of the covariate experienced by each species. Truncated trendlines occur for *B. mixtus* because its shorter foraging range limits the maximum accessible habitat area. Shading indicates 95% credible intervals: a solid trendline and coloured shading indicated that the 95% credible interval for an association excluded zero (“strong support”); a dashed trendline and coloured shading indicates that the 90% credible interval for an association excluded zero (“moderate support”). Grey shading indicates an assocation with weak or no support. Inset figures show posteriors for the average marginal effect (AME) of each predictor. AMEs are computed for a one standard deviation increase in each predictor and are based on predictions for one hour of survey time. Vertical red dashed lines in the inset indicate zero. *P*(Θ): posterior density.

Results of worker abundance models were qualitatively similar to the colony abundance models, although the support for a negative association between berry crop area and *B. impatiens* worker abundance was slightly stronger, and the support a positive association between seminatural area and *B. mixtus* worker abundance was slightly weaker. Full results can be found in Fig. A5 and Table A2. Worker and colony counts were highly correlated for both *B. mixtus* (*r*^2^ = 0.988) and *B. impatiens* (*r*^2^ = 0.989).

### 3.3 Queen abundance

Queens of both species were more abundant at transect-surveys with higher floral abundance; while model support was strong for *B. mixtus*, it was only moderate for *B. impatiens*. A one standard deviation increase (i.e., 1.15 unit increase) in log-scaled floral abundance was associated with an increase of 0.42 (95% CrI: [0.15, 0.77]) *B. mixtus* queens and 0.22 (95% CrI: -0.03-0.53) *B. impatiens* queens per hour (Fig. 3 A-B).

**Figure 3:**
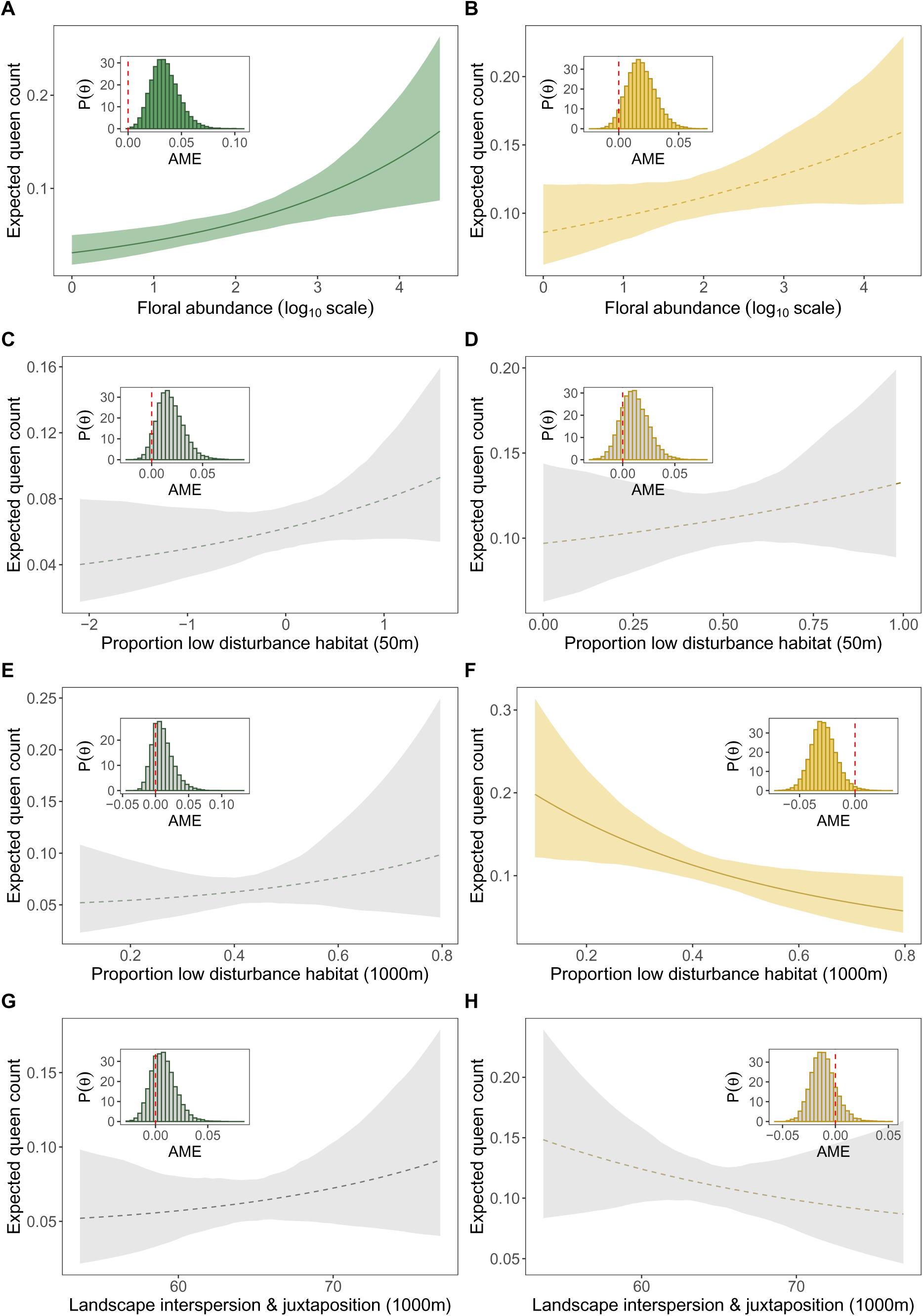
Queen abundance for *B. mixtus* (lefthand column, green) and *B. impatiens* (righthand column, yellow) in relation to floral abundance (A-B), proportion low disturbance habitat in a 50m buffer (C-D), proportion low disturbance habitat in a 1km buffer (E-F) and landscape interspersion and juxtaposition within a 1km buffer (G-H). Shading indicates 95% credible intervals: a solid trendline and coloured shading indicated that the 95% credible interval for an association excluded zero (“strong support”); a dashed trendline and coloured shading indicates that the 90% credible interval for an association excluded zero (“moderate support”); grey shading indicates an assocation with weak or no support. Inset figures show posteriors for the average marginal effect (AME) of a one standard deviation increase in each predictor, computed for one hour of survey time. *P*(Θ): posterior density.

There was no significant effect of local-scale low-disturbance habitat (Fig. 3 C-D) or landscape interspersion and juxtaposition index (Fig. 3G-H) on queen abundance of either species. *B. mixtus* queen abundance was not associated with landscape-scale low-disturbance habitat, but *B. impatiens* queen abundance decreased by an average of 0.34 (95% CrI: [0.05, 0.60]) individuals per hour when landscape-scale low disturbance habitat increased by one standard deviation (i.e., 17.5%) (Fig. 3 E-F).

### 3.4 Queen behaviour

The probability of queens engaging in nest searching rather than foraging was not strongly associated with local floral abundance for either species (Fig. 4 A-B). There was moderate support that *B. mixtus* was more likely to nest-search in blueberry fields (ΔP(nest searching) = 0.2 (95% CrI: [−0.03, 0.45])) and field margins (ΔP(nest searching) = 0.19 (95% CrI: [−0.01, 0.39])) in comparison to roadsides. We did not find evidence for an effect of local habitat type (roadside, garden, seminatural, field edge, blueberry) on *B. impatiens* queen behaviour (Fig. 4C).

**Figure 4:**
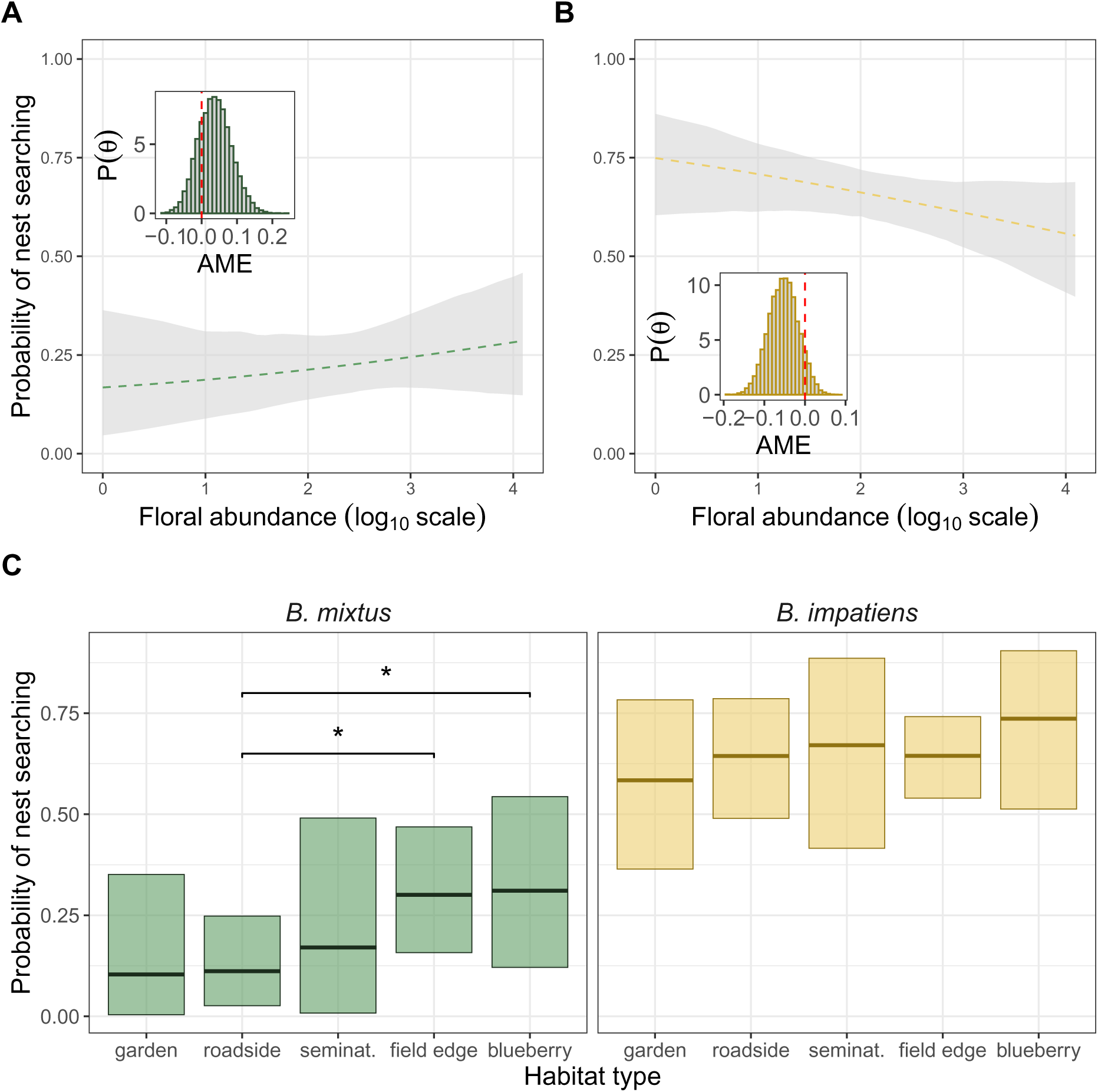
Queen behaviour (1 = nest searching, 0 = foraging) in relation to local transect characteristics for *B. mixtus* (green) and *B. impatiens* (yellow). (A-B) Predicted probability of nest searching as a function of floral abundance; shading indicates 95% credible interval. Inset figures show posteriors for the average marginal effect (AME) of a one standard deviation increase in *log*_10_ floral abundance. Grey shading and a dashed trendline are utilized when the 95% credible interval for the AME contains zero. (C) Probability of nest searching for each species at transects within each habitat type; shading indicates 95% credible interval, a single star (*) indicates moderate support for a difference in nest searching probability between factor levels. *P*(Θ): posterior density.

**Figure 5:**
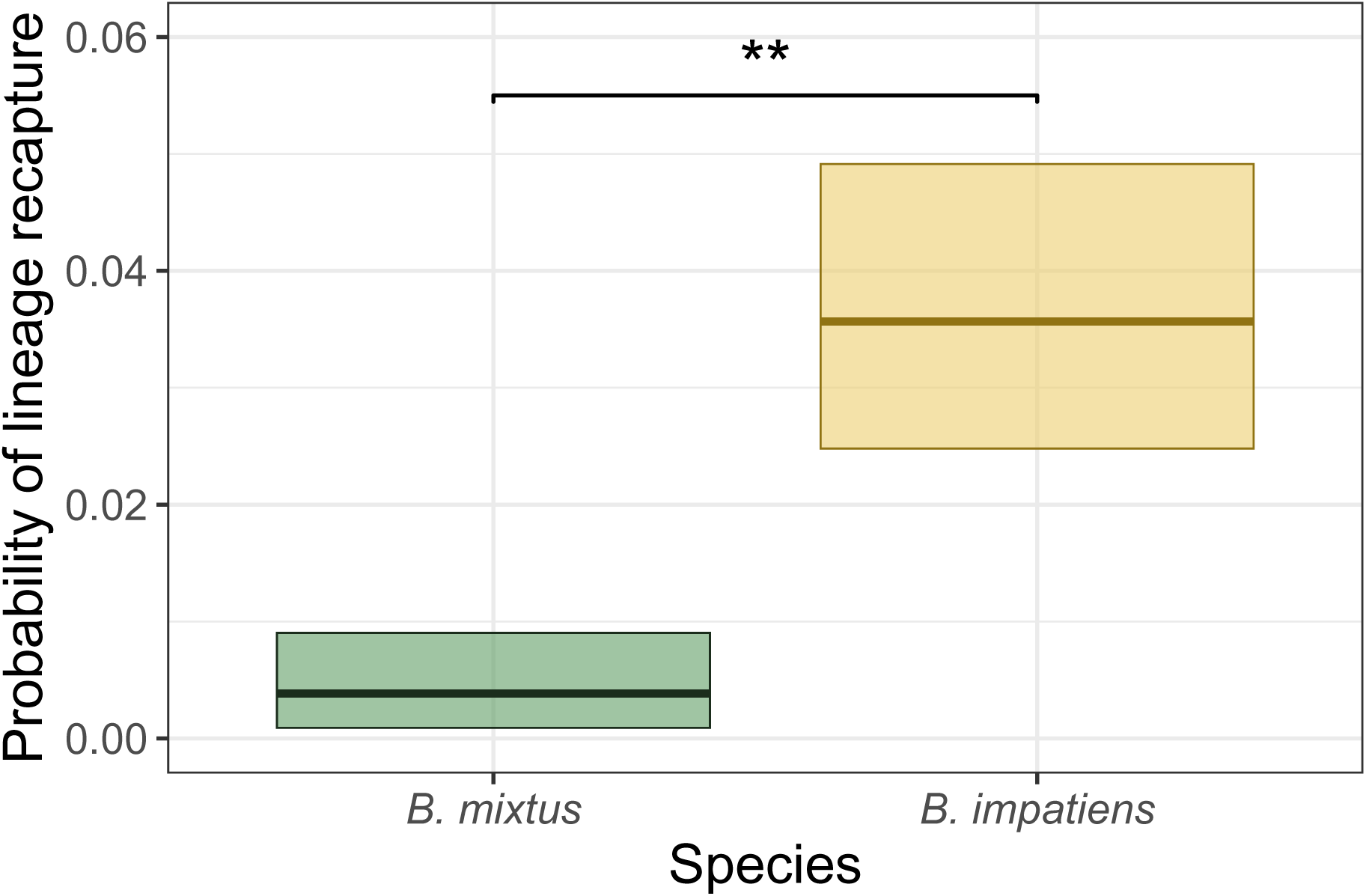
Rates of lineage recapture between 2022 colonies and 2023 spring queens for *B. mixtus* and *B. impatiens*. Shading indicates 95% credible interval. Two stars (**) indicate strong support for a different in probability between species.

The strongest determinant of nest-searching behaviour was species identity: *B. impatiens* had a marginal expected probability of nest searching of 0.65 (95% CrI: [0.59, 0.71]) while *B. mixtus* had a probability of nest searching of 0.22 (95% CrI: [0.14, 0.31]) (*P_impatiens_* − *P_mixtus_* = 0.43, 95% CrI: [0.32, 0.53]) (Fig. 4C). Additional analyses showed that species identity was an important predictor of nest-searching behaviour across the entire bumblebee community, with species-level probability of nest-searching varying from 0.07 (95% CrI: [0.0022, 0.23]) for *B. flavifrons* to 0.73 (95% CrI: [0.59, 0.84]) for *B. vosnesenskii* (Fig. A6).

### 3.5 Lineage survival

In the spring of 2023, we re-observed 33 colony lineages (out of 1,023 initially detected in 2022) for *B. impatiens* and 3 colony lineages (out of 673 initially detected in 2022) for *B. mixtus*. The marginal probability of lineage survival was greater for *B. impatiens* (0.036, 95% CrI: [0.025, 0.049]) than for *B. mixtus* (0.0038, 95% CrI: [0.00082, 0.0091]) (*P_impatiens_* − *P_mixtus_* = 0.032, 95% CrI: [0.019, 0.046]). One lineage of *B. impatiens* was observed in all three time periods (2022 workers, 2023 queens, 2023 workers).

## 4 Discussion

4.1 Landscape composition differentially impacts the abundance of *B. mixtus* and

### B. impatiens

As predicted, we observed different spatial patterns of colony, worker, and queen abundance for *B. mixtus* and *B. impatiens*. Throughout the region and in both years, *B. impatiens* colony and queen abundances were higher than those of *B. mixtus*. The two species had different responses to seminatural habitat, with *B. mixtus* colony abundance trending upwards and *B. impatiens* abundance trending downwards in response to increasing seminatural habitat area. This result is unsurprising for two reasons: first, much of the seminatural habitat in this region consists of forest fragments, which are mostly closed-canopy coniferous forest that likely provide unfamiliar foraging/nesting habitat for *B. impatiens* given its history in the deciduous forests/open habitat of eastern North America. In contrast, *B. mixtus* and other early-emerging native bumble bees are highly attracted to understory shrubs such as Salmonberry (*Rubus spectablis*) and Common Snowberry (*Symphoricarpos albus*) which can be found in forest fragments. In general, populations of *B. impatiens* in this region have a stronger preference for introduced plant species in comparison to native bumble bees (Platsko et al., 2025). Second, in its native range *B. impatiens* is found at higher abundance in human-modified versus natural landscapes (Gratton et al., 2023), suggesting that it may generally thrive under the biotic and abiotic conditions produced by agricultural intensification. The association of *B. impatiens* with human-modified landscapes is further supported by a higher abundance of *B. impatiens* colonies in landscapes with more residential garden area. Due to the later phenology and wide diet breadth of *B. impatiens* (Platsko et al., 2025), we hypothesize that this species may benefit from late-season horticultural plants available in gardens. As a result, while maintaining seminatural habitat fragments may be key for supporting *B. mixtus* and potentially other native *Bombus*, loss of these habitats or increased suburbanization may support the spread and proliferation of *B. impatiens*.

Both species showed positive trends in colony abundance in relation to field margin (edge) area, which were moderately supported in *B. impatiens* but only weakly supported for *B. mixtus*. Other recent studies have also highlighted the importance of field edge density within agricultural landscapes for increasing biodiversity (Fahrig et al., 2015; Sirami et al., 2019), although in our system species-specific effects may depend on the floral resources offered by field margins: if these are mainly arable weeds, an introduced generalist like *B. impatiens* may benefit disproportionately from these habitats. In support of this hypothesis, 88.2% (4422/5015) of all floral observations in edge habitats (field margins and roadsides) were exotic or crop plants; we further observed that 90.3% (2984/3304) of all *B. impatiens* visits were to non-native flowers, in comparison to 73.6% (1457/1980) for *B. mixtus*.

Both species exhibited negative trends in response to increased area of intensively managed berry-crops (blueberry and cranberry); the effect size of this relationship was larger and more clearly supported for *B. mixtus* than for *B. impatiens*. We included berry crops in our model as potential foraging habitat because they are mass-flowering, but there are several reasons why these crops may instead limit *B. mixtus* populations. Berry crops in this region are highly dominated by blueberry, a pollinator-dependent crop which is managed intensively with fungicide, herbicide, and insecticide applications to suppress pest populations and weed growth—as a result, fields are typically resource-poor outside of the bloom period (May-June) and may expose foraging or nesting bumble bees to high pesticide risk. In addition, blueberry itself provides very little nectar and its pollen is of low nutritional quality for foraging bumble bees (Toshack & Elle, 2019). The weaker response of *B. impatiens* to these intensively managed crops could result from (i) lower exposure to pesticides via foraging, as its later phenology leads workers to emerge after the blueberry bloom, (ii) higher tolerance to pesticides due to larger body size, or (iii) greater ability to utilize nesting sites in blueberry fields.

Lastly, we found that while the quantity of low-disturbance habitat within 50 meters of each transect had little effect on the abundance of queens for either species, *B. impatiens* queens were observed in lower numbers when landscape-scale low-disturbance habitat proportion was higher. We hypothesize that the observed pattern arises through concentration of nest searching queens. In general, we expect queens to be evenly distributed across potentially suitable sites; when the proportion of suitable habitat in a landscape is low, nest-searching queens must concentrate themselves on the few undisturbed habitat elements available. Because the majority of our surveys took place in areas which were presumably suitable for nesting (field margins, gardens, roadsides, etc.), this concentration could explain the increase in queen observations when landscape-scale nesting habitat is rare. The fact that we do not observe a similar trend for *B. mixtus* queens could mean that agricultural nest-site limitation does not drive patterns of space use for this species, potentially because it finds agricultural landscapes less suitable for nesting (discussed below). Alternatively, *B. mixtus* may have more restrictive microhabitat requirements, such that our classification of suitable/unsuitable nesting habitat is not informative.

Together these results support the hypothesis that *B. impatiens*, a highly successful and expanding introduced species, is more strongly associated with agricultural and residential land use than its native congener, and is relatively insensitive to intensive crop management. In contrast to *B. mixtus*, its abundance is lower in landscapes with more seminatural habitat and higher in landscapes with more residential gardens, suggesting that either the two species receive different utility from these habitat types or that *B. impatiens* has so far failed to colonize sites where native bumble bees are present in higher numbers.

### 4.2 Behavioural differences may underpin patterns of abundance

We show that the mean and maximum foraging distance of *B. impatiens* are larger than those of *B. mixtus*; our spatial models confirm that there is a significant difference in the foraging scales of the two species. This result is perhaps unsurprising given the larger body size of *B. impatiens* (Williams et al., 2014), although body size is not necessarily a good predictor of foraging range within the genus *Bombus* (Redhead et al., 2016). Differences in foraging range may also reflect subtle differences in life history traits, such as colony size, which is hypothesized to be linked to foraging distance when food availability is limited (Grüter & Hayes, 2022). While colony size is not well studied for *B. mixtus*, *B. impatiens* nests unearthed in northern Washington hosted up to 1000 brood cells (JBUK, personal observation), which is on the larger side for *Bombus*.

Ultimately, we hypothesize that the larger foraging range of *B. impatiens* may contribute to its success in human-modified landscapes, where floral resources can be highly abundant but temporally transient. Walther-Hellwig and Frankl (2000) showed than in the UK, species with larger foraging ranges were more likely to concentrate on mass flowering crops, while species with restricted foraging ranges relied on close proximity between nesting and foraging habitat. Similarly, *B. impatiens* may be less sensitive to intensively managed berry crops (and their post-bloom resource dearth) if greater plasticity in its foraging range allows it to travel further between nesting and foraging habitat.

In addition to differences in foraging range, *B. mixtus* and *B. impatiens* exhibit marked differences in their apparent allocation of time spent nest searching versus foraging at our sites, with nearly 65% of all *B. impatiens* queens displaying nest searching behaviour, in comparison to around 20% of *B. mixtus* queens. These behaviours align with our observation that *B. mixtus* queen abundance is most strongly correlated with floral abundance while *B. impatiens* queen abundance is also shaped by landscape-scale nesting habitat availability. We posit that these differences may reflect the preferred nesting locations/substrates of the two species; while *B. impatiens* is a primarily underground nester, *B. mixtus* often creates surface nests (Williams et al., 2014). With this context, our analysis suggests that habitats which are suitable nesting habitat for *B. impatiens* (e.g., areas without tillage such as blueberry fields, roadsides or field margins) are only partially suitable for *B. mixtus*, perhaps because these areas still receive surface-level disturbances via mowing or vehicle use. This hypothesis is further supported by the moderately supported differences in *B. mixtus* nest searching probability across habitat types, which may suggest greater selectivity in nesting locations in comparison to *B. impatiens*. Given the high disturbance rates of many agricultural landcover types, it is possible that *B. mixtus* queens primarily use agricultural landscapes for foraging, rather than nest searching/nesting. Alternatively, *B. mixtus* and other surface-nesting species may preferentially nest-search in more vegetated areas (e.g., in tall grass or shrubs, or in the interior of a hedgerow rather than along its edge), decreasing their detectability in comparison to *B. impatiens*. Both hypotheses are supported by our observations of other *Bombus spp.* queens; the three species most commonly observed nest searching rather than foraging were *B. huntii*, *B. impatiens*, and *B. vosnesenskii*—the only primarily belowground-nesting species we observed (Williams et al., 2014). We do not believe these patterns are attributable to sampling timing; our queen observations began in mid-March, as soon as the temperature rose above 10^◦^C. Below this temperature bumblebee queens are not typically active in our region; furthermore, if the first stages of queen emergence were missed (during which queens typically forage for resources to regain energy post-hibernation (Williams et al., 2014)), it would result in an over-estimation of nest-searching behaviour for early emerging species (*B. melanopygus, B. flavifrons, B. mixtus*), a pattern which does not align with our observations.

### 4.3 Colony lineage recapture is higher for *B. impatiens* than for *B. mixtus*

By assigning spring queens to their natal colonies, we were able to assess and compare the lineage recapture rates of both species. We found that the probability of lineage recapture was nearly 10 times higher for *B. impatiens* (3.6%) than for *B. mixtus* (0.38%). This result is consistent with the hypothesis that *B. impatiens* has higher rates of colony survival and/or reproductive success in agricultural landscapes than *B. mixtus*, which may be a key driver of its expansion in this region. However, it is important to note that failure to recapture a lineage could be due to a variety of factors other than colony failure, such as dispersal of reproductive offspring away from the natal site. Given our observation that *B. mixtus* has a smaller foraging range than *B. impatiens*, we hypothesize that the average dispersal distance of *B. mixtus* is also smaller. This should produce a *higher* recapture rate for this species, all else equal, and suggests that the direction of the disparity we observed is due to real differences in reproductive success. However, different environmental cues or intrinsic behaviours are likely to guide foraging and dispersal (Mola & Williams, 2025), so comparative studies of queen dispersal would be necessary to validate this hypothesis.

### 4.4 Conclusions and future directions

Globally, landscapes are undergoing modification for agriculture and residential development, creating mosaics of agricultural, (sub)urban and seminatural landcover, which experience different degrees of disturbance due to human activities. Our results suggest that an introduced pollinator (*B. impatiens*) is more strongly associated with highly human-modified landscapes than its native relative (*B. mixtus*); higher lineage recapture rates of the introduced species seem to indicate greater reproductive success in agro-natural mosaics, a finding which is supported by its rapid rise in abundance in this region. Our results align with previous trait-based time-series analyses which suggest that habitat generalists and species with strong affinity for human-modified habitats are more likely to persist following anthropogenic change (Fisogni et al., 2025). Our research provides novel evidence suggesting that behaviours such as foraging range and nest establishment may underpin this trait-filtering, and that these behaviours are sufficient to explain differential population trends between closely related (congeneric) species.

Our findings also suggest that loss of seminatural habitat and increased suburbanization/agricultural intensification may facilitate the expansion of highly successful species like *B. impatiens*, creating spatial patterns of abundance which differ from those of closely related native species. This possibility brings into question the continued use of *B. impatiens* and species like it as model organisms for studying population responses to land use change; future work should assess colony survival and reproductive rates of *B. impatiens* across a gradient of land use intensification in this region and in its native range, ideally in contrast to a less successful native relative. Understanding how land use change impacts the demography of introduced and expanding species is critical for predicting and managing their spread and forecasting future provisioning of ecosystem services.

## 5 Declaration of generative AI and AI-assisted technologies in the manuscript preparation process

During the preparation of this work the author(s) used Claude to debug R code and to perform spot-edits for grammar and clarity. All code was tested prior to application to data, and suggested text edits (e.g., rephrasing of sentences) were incorporated manually without changes to the content/terminology of the original draft.

## Author contributions

JBM and CK conceived the ideas, all authors contributed to methodological design; JBM, NC, and TTK collected the data; JBM analysed the data and led the writing of the manuscript. All authors contributed critically to the drafts and gave final approval for publication.

## Acknowledgements

The authors gratefully acknowledge Charles Margossian and Victor Van der Meersch for many helpful conversations on modelling foraging distance, and Michael Whitlock for advice and feedback on population genetic analyses. We thank Hazel Barthel and Elinor Sisk for their invaluable assistance with field data collections. This study would not have been possible without the support of landowners/managers throughout the Lower Fraser Valley. The research was supported by the Natural Sciences and Engineering Research Council of Canada (NSERC) [ALLRP 570736-2021 and RGPIN-2020-05759], the Canada Foundation for Innovation / John R. Evans Leaders Fund (40541) in collaboration with British Columbia Knowledge Development Fund (BCKDF), and the AgriScience Program under Agriculture and Agri-Food Canada’s Sustainable Canadian Agricultural Partnership as part of Organic Science Cluster 4 (to CK), as well as USDA grant 2022-67013-36286 (to JBUK). JBM was supported by NSERC Vanier CGS 01353 - 000.

## Data Availability

Version-controlled code necessary to reproduce the analyses can be found at https://github.com/jmelanson98/fv_landscapeforaging. Full datasets will be shared to the same repository and archived on Dryad following acceptance.

