## Appendix for "Agricultural intensification favours an introduced bumble bee over its native congener through differences in foraging range, habitat association, and lineage continuity"

### A1 Additional Tables and Figures

Table A1: Summary of coefficients for colony abundance models (conditional coefficients on the linear-predictor scale). All "area" predictors are distance-weighted areas, computed as described in the Methods section of the main text. Predictors were centered and scaled prior to model fitting. IJI = interspersion and juxtaposition index. ESS = Effective Sample Size.

| Parameter | Estimate | SE | $CI_{2.5\%}$ | $CI_{97.5\%}$ | $ESS_{bulk}$ | $ESS_{tail}$ | $\hat{R}$ |
|---|---|---|---|---|---|---|---|
| *B. impatiens* intercept | -2.32 | 0.18 | -2.67 | -1.96 | 5683 | 5751 | 1.00 |
| *B. mixtus* intercept | -2.71 | 0.18 | -3.06 | -2.34 | 5902 | 5882 | 1.00 |
| Blueberry area | -0.14 | 0.09 | -0.31 | 0.03 | 8528 | 10685 | 1.00 |
| Residential garden area | 0.14 | 0.07 | 0.00 | 0.29 | 8288 | 10255 | 1.00 |
| Field edge area | 0.13 | 0.08 | -0.02 | 0.28 | 8448 | 10908 | 1.00 |
| Fallow area | 0.01 | 0.07 | -0.11 | 0.15 | 9474 | 11296 | 1.00 |
| Seminatural area | -0.11 | 0.07 | -0.24 | 0.02 | 9354 | 10875 | 1.00 |
| IJI | -0.04 | 0.09 | -0.22 | 0.13 | 8966 | 11744 | 1.00 |
| Year (2023) | -0.12 | 0.11 | -0.33 | 0.09 | 13908 | 12394 | 1.00 |
| Floral abundance | 0.22 | 0.08 | 0.07 | 0.37 | 11097 | 12517 | 1.00 |
| Blueberry area : *B. mixtus* | -0.22 | 0.10 | -0.41 | -0.03 | 16504 | 13289 | 1.00 |
| Residential garden area : *B. mixtus* | -0.12 | 0.08 | -0.29 | 0.04 | 13624 | 12286 | 1.00 |
| Field edge area : *B. mixtus* | -0.06 | 0.08 | -0.21 | 0.10 | 15745 | 12033 | 1.00 |
| Fallow area : *B. mixtus* | 0.01 | 0.07 | -0.13 | 0.16 | 17945 | 12545 | 1.00 |
| Seminatural area : *B. mixtus* | 0.23 | 0.07 | 0.09 | 0.37 | 14748 | 12543 | 1.00 |
| IJI : *B. mixtus* | 0.02 | 0.09 | -0.17 | 0.20 | 13186 | 13311 | 1.00 |
| Year (2023) : *B. mixtus* | -0.39 | 0.15 | -0.69 | -0.09 | 15023 | 12457 | 1.00 |
| Floral abundance : *B. mixtus* | 0.23 | 0.10 | 0.04 | 0.42 | 16021 | 12853 | 1.00 |

Table A2: Summary of coefficients for worker abundance models (conditional coefficients on the linear-predictor scale). All "area" predictors are distance-weighted areas, computed as described in the Methods section of the main text. Predictors were centred and scaled prior to model fitting. IJI = interspersion and juxtaposition index. ESS = Effective Sample Size.

| Parameter | Estimate | SE | $CI_{2.5\%}$ | $CI_{97.5\%}$ | $ESS_{bulk}$ | $ESS_{tail}$ | $\hat{R}$ |
|---|---|---|---|---|---|---|---|
| *B. impatiens* intercept | -2.28 | 0.19 | -2.65 | -1.90 | 5397 | 6318 | 1.00 |
| *B. mixtus* intercept | -2.67 | 0.19 | -3.05 | -2.29 | 5430 | 5944 | 1.00 |
| Blueberry area | -0.16 | 0.09 | -0.35 | 0.02 | 9838 | 11633 | 1.00 |
| Residential garden area | 0.14 | 0.08 | 0.00 | 0.29 | 10826 | 11324 | 1.00 |
| Field edge area | 0.15 | 0.08 | -0.01 | 0.31 | 10398 | 11435 | 1.00 |
| Fallow area | 0.02 | 0.07 | -0.12 | 0.15 | 11003 | 11589 | 1.00 |
| Seminatural area | -0.11 | 0.07 | -0.25 | 0.03 | 10939 | 12592 | 1.00 |
| IJI | -0.05 | 0.09 | -0.23 | 0.14 | 9769 | 11315 | 1.00 |
| Year (2023) | -0.08 | 0.11 | -0.30 | 0.14 | 14121 | 12773 | 1.00 |
| Floral abundance | 0.21 | 0.08 | 0.06 | 0.37 | 12582 | 10957 | 1.00 |
| Blueberry area : *B. mixtus* | -0.22 | 0.10 | -0.42 | -0.03 | 17862 | 12457 | 1.00 |
| Residential garden area : *B. mixtus* | -0.12 | 0.09 | -0.29 | 0.04 | 17685 | 12825 | 1.00 |
| Field edge area : *B. mixtus* | -0.07 | 0.08 | -0.23 | 0.09 | 17940 | 13225 | 1.00 |
| Fallow area : *B. mixtus* | 0.01 | 0.08 | -0.14 | 0.16 | 20702 | 12900 | 1.00 |
| Seminatural area : *B. mixtus* | 0.22 | 0.07 | 0.08 | 0.37 | 17870 | 12026 | 1.00 |
| IJI : *B. mixtus* | 0.04 | 0.10 | -0.16 | 0.23 | 14762 | 12397 | 1.00 |
| Year (2023) : *B. mixtus* | -0.43 | 0.16 | -0.74 | -0.13 | 15494 | 12475 | 1.00 |
| Floral abundance : *B. mixtus* | 0.23 | 0.10 | 0.04 | 0.43 | 16317 | 12280 | 1.00 |

Table A3: Summary of coefficients for *B. mixtus* queen abundance models (conditional coefficients on the linear-predictor scale). Continuous predictors were scaled and centered prior to model fitting. IJI = interspersion and juxtaposition index. ESS = Effective Sample Size.

| Parameter | Estimate | SE | $CI_{2.5\%}$ | $CI_{97.5\%}$ | $ESS_{bulk}$ | $ESS_{tail}$ | $\hat{R}$ |
|---|---|---|---|---|---|---|---|
| Intercept | -2.63 | 0.49 | -3.61 | -1.67 | 4583 | 7200 | 1.00 |
| Low disturbance area (50m) | 0.24 | 0.15 | -0.05 | 0.55 | 13111 | 12201 | 1.00 |
| Low disturbance area (1000m) | 0.14 | 0.21 | -0.26 | 0.57 | 10258 | 11795 | 1.00 |
| IJI (1000m) | 0.10 | 0.17 | -0.23 | 0.42 | 13381 | 12496 | 1.00 |
| Floral abundance | 0.43 | 0.13 | 0.18 | 0.67 | 19661 | 13622 | 1.00 |
| Julian date | -2.12 | 0.34 | -2.84 | -1.50 | 11242 | 10322 | 1.00 |
| Julian date$^2$ | -1.47 | 0.25 | -1.98 | -1.01 | 12381 | 10932 | 1.00 |

Table A4: Summary of coefficients for *B. impatiens* queen abundance models (conditional coefficients on the linear-predictor scale). Continuous predictors were scaled and centred prior to model fitting. IJI = interspersion and juxtaposition index. ESS = Effective Sample Size.

| Parameter | Estimate | SE | $CI_{2.5\%}$ | $CI_{97.5\%}$ | $ESS_{bulk}$ | $ESS_{tail}$ | $\hat{R}$ |
|---|---|---|---|---|---|---|---|
| Intercept | -1.41 | 0.31 | -2.01 | -0.81 | 4482 | 7694 | 1.00 |
| Low disturbance area (50m) | 0.09 | 0.11 | -0.12 | 0.30 | 10676 | 11345 | 1.00 |
| Low disturbance area (1000m) | -0.30 | 0.14 | -0.58 | -0.04 | 9902 | 11219 | 1.00 |
| IJI (1000m) | -0.11 | 0.12 | -0.34 | 0.12 | 9775 | 10055 | 1.00 |
| Floral abundance | 0.15 | 0.09 | -0.02 | 0.33 | 16804 | 12795 | 1.00 |
| Julian date | -1.17 | 0.18 | -1.54 | -0.84 | 12688 | 11549 | 1.00 |
| Julian date$^2$ | -1.49 | 0.17 | -1.82 | -1.18 | 12962 | 11411 | 1.00 |

Table A5: Summary of coefficients for *B. mixtus* and *B. impatiens* queen behaviour models (conditional coefficients on the linear-predictor scale). Continuous predictors were centred and scaled prior to model fitting. ESS = Effective Sample Size.

| Parameter | Estimate | SE | $CI_{2.5\%}$ | $CI_{97.5\%}$ | $ESS_{bulk}$ | $ESS_{tail}$ | $\hat{R}$ |
|---|---|---|---|---|---|---|---|
| *B. impatiens* intercept | 2.01 | 0.85 | 0.44 | 3.79 | 4832 | 6334 | 1.00 |
| *B. mixtus* intercept | -0.70 | 1.04 | -2.84 | 1.25 | 6391 | 8190 | 1.00 |
| *B. impatiens* : field margin | -0.68 | 0.85 | -2.39 | 0.95 | 5007 | 6928 | 1.00 |
| *B. mixtus* : field margin | -0.02 | 0.98 | -1.89 | 1.96 | 7130 | 9252 | 1.00 |
| *B. impatiens* : garden | -1.00 | 1.06 | -3.09 | 1.10 | 6009 | 8055 | 1.00 |
| *B. mixtus* : garden | -2.26 | 1.70 | -6.00 | 0.75 | 10939 | 10905 | 1.00 |
| *B. impatiens* : roadside | -0.67 | 0.94 | -2.57 | 1.18 | 5178 | 6543 | 1.00 |
| *B. mixtus* : roadside | -1.78 | 1.11 | -4.05 | 0.33 | 8999 | 11217 | 1.00 |
| *B. impatiens* : seminatural | -0.46 | 1.14 | -2.69 | 1.79 | 6849 | 9547 | 1.00 |
| *B. mixtus* : seminatural | -1.43 | 1.69 | -5.10 | 1.53 | 12244 | 10545 | 1.00 |
| *B. impatiens* : floral abundance | -0.32 | 0.23 | -0.77 | 0.11 | 14782 | 12144 | 1.00 |
| *B. mixtus* : floral abundance | 0.31 | 0.39 | -0.42 | 1.11 | 9830 | 11870 | 1.00 |
| *B. impatiens* : Julian date | -0.50 | 0.57 | -1.63 | 0.60 | 11729 | 10825 | 1.00 |
| *B. mixtus* : Julian date | -0.54 | 1.58 | -3.88 | 2.40 | 9055 | 8729 | 1.00 |
| *B. impatiens* : Julian date$^2$ | -1.31 | 0.50 | -2.36 | -0.37 | 10853 | 10771 | 1.00 |
| *B. mixtus* : Julian date$^2$ | -1.73 | 1.28 | -4.48 | 0.54 | 10734 | 9831 | 1.00 |

Table A6: Summary of coefficients for models of queen behaviour across all species (conditional coefficients on the linear-predictor scale). Continous predictors were centred and scaled prior to model fitting. ESS = Effective Sample Size.

| Parameter | Estimate | SE | $CI_{2.5\%}$ | $CI_{97.5\%}$ | $ESS_{bulk}$ | $ESS_{tail}$ | $\hat{R}$ |
|---|---|---|---|---|---|---|---|
| Intercept (*B. californicus* / blueberry) | -1.97 | 1.01 | -4.08 | -0.09 | 3970 | 4720 | 1.00 |
| Field margin habitat type | -0.06 | 0.60 | -1.20 | 1.16 | 4548 | 5204 | 1.00 |
| Garden habitat type | -0.98 | 0.77 | -2.49 | 0.53 | 5573 | 6650 | 1.00 |
| Roadside habitat type | -0.57 | 0.62 | -1.79 | 0.68 | 4755 | 6643 | 1.00 |
| Seminatural habitat type | -0.29 | 0.79 | -1.83 | 1.28 | 5820 | 8438 | 1.00 |
| Floral abundance | -0.30 | 0.18 | -0.64 | 0.05 | 8920 | 10101 | 1.00 |
| *B. flavifrons* | -1.08 | 1.58 | -4.50 | 1.72 | 6805 | 8973 | 1.00 |
| *B. huntii* | 3.42 | 1.57 | 0.51 | 6.75 | 6150 | 7098 | 1.00 |
| *B. impatiens* | 3.58 | 0.84 | 2.10 | 5.44 | 3823 | 4219 | 1.00 |
| *B. melanopygus* | 3.17 | 1.24 | 0.80 | 5.66 | 5723 | 6554 | 1.00 |
| *B. mixtus* | 0.98 | 0.87 | -0.60 | 2.84 | 3964 | 4876 | 1.00 |
| *B. nevadensis* | -709.17 | 998.33 | -3396.60 | -12.44 | 1853 | 831 | 1.00 |
| *B. rufocinctus* | 1.23 | 1.08 | -0.83 | 3.42 | 5302 | 5480 | 1.00 |
| *B. sitkensis* | -371.99 | 607.15 | -1706.06 | -7.12 | 1646 | 708 | 1.00 |
| *B. vosnesenskii* | 4.26 | 0.95 | 2.58 | 6.29 | 4085 | 4939 | 1.00 |
| Julian date | -0.96 | 0.43 | -1.83 | -0.16 | 7645 | 8727 | 1.00 |
| Julian date$^2$ | -1.52 | 0.38 | -2.31 | -0.80 | 8350 | 9264 | 1.00 |

Table A7: Summary of coefficients for lineage survival models (conditional coefficients on the linear-predictor scale). ESS = Effective Sample Size.

| Parameter | Estimate | SE | $CI_{2.5\%}$ | $CI_{97.5\%}$ | $ESS_{bulk}$ | $ESS_{tail}$ | $\hat{R}$ |
|---|---|---|---|---|---|---|---|
| *B. impatiens* | -3.49 | 0.57 | -4.70 | -2.41 | 1531 | 1842 | 1.00 |
| *B. mixtus* | -5.93 | 0.83 | -7.76 | -4.45 | 2311 | 2720 | 1.00 |

Table A8: Results of Moran's I test at two spatial scales for all model residuals. We tested for positive spatial autocorrelation in model residuals, i.e. whether residuals of spatially proximate points were more similar than expected under spatial randomness. While p-values close to 1 could be indicative of negative spatial autocorrelation, we note that the magnitude of Moran's I statistic for these values (Moran's I < 0.1) is likely too small to hold much biological relevance. E[I]: expectation of Moran's I.

| Model (Spatial Scale) | Moran's $I$ | $E[I]$ | Variance | $p$-value |
|---|---|---|---|---|
| Colony abundance (500m) | -3.481e-02 | -1.056e-03 | 8.248e-05 | 9.999e-01 |
| Colony abundance (1000m) | -8.794e-03 | -1.056e-03 | 2.444e-05 | 9.412e-01 |
| Worker abundance (500m) | -3.410e-02 | -1.056e-03 | 8.311e-05 | 9.999e-01 |
| Worker abundance (1000m) | -9.044e-03 | -1.056e-03 | 2.462e-05 | 9.463e-01 |
| *B. mixtus* queen abundance (500m) | -3.410e-02 | -1.056e-03 | 8.311e-05 | 9.999e-01 |
| *B. impatiens* queen abundance (1000m) | -8.794e-03 | -1.056e-03 | 2.444e-05 | 9.412e-01 |
| Queen behaviour (500m) | -3.410e-02 | -1.056e-03 | 8.311e-05 | 9.999e-01 |
| Queen behaviour (1000m) | -8.794e-03 | -1.056e-03 | 2.444e-05 | 9.412e-01 |
| Queen behaviour (all species) (500m) | -3.410e-02 | -1.056e-03 | 8.311e-05 | 9.999e-01 |
| Queen behaviour (all species) (1000m) | -8.794e-03 | -1.056e-03 | 2.444e-05 | 9.412e-01 |

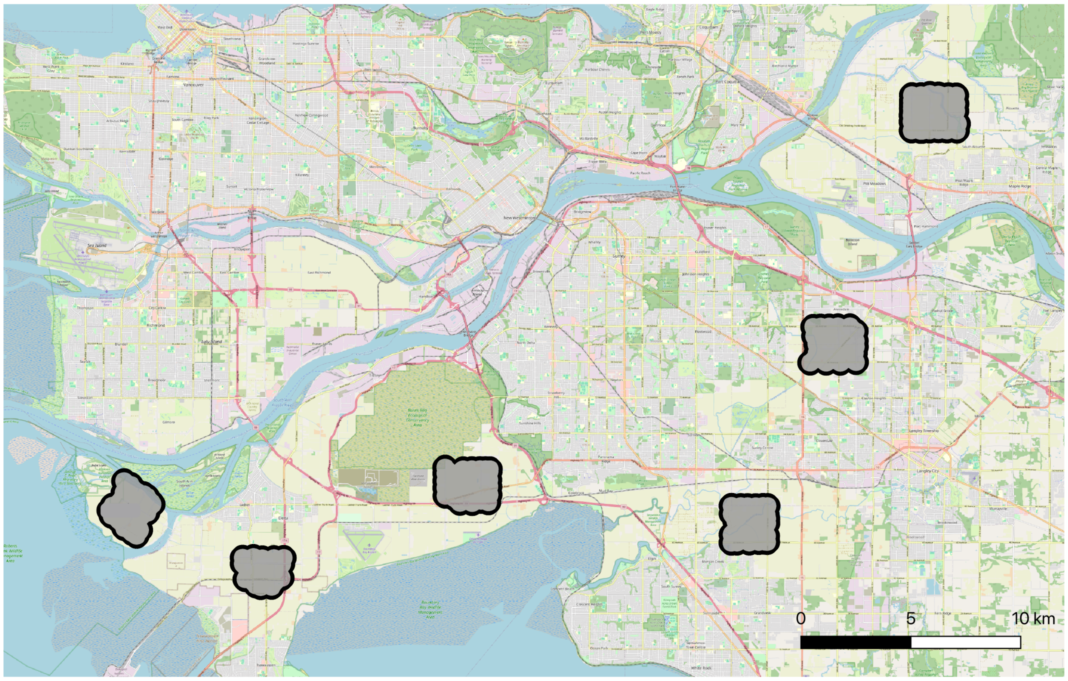


Figure A1: Our study region, located near Vancouver, British Columbia, Canada (city in top left corner). The six replicate landscapes are shown in grey, with black outline.

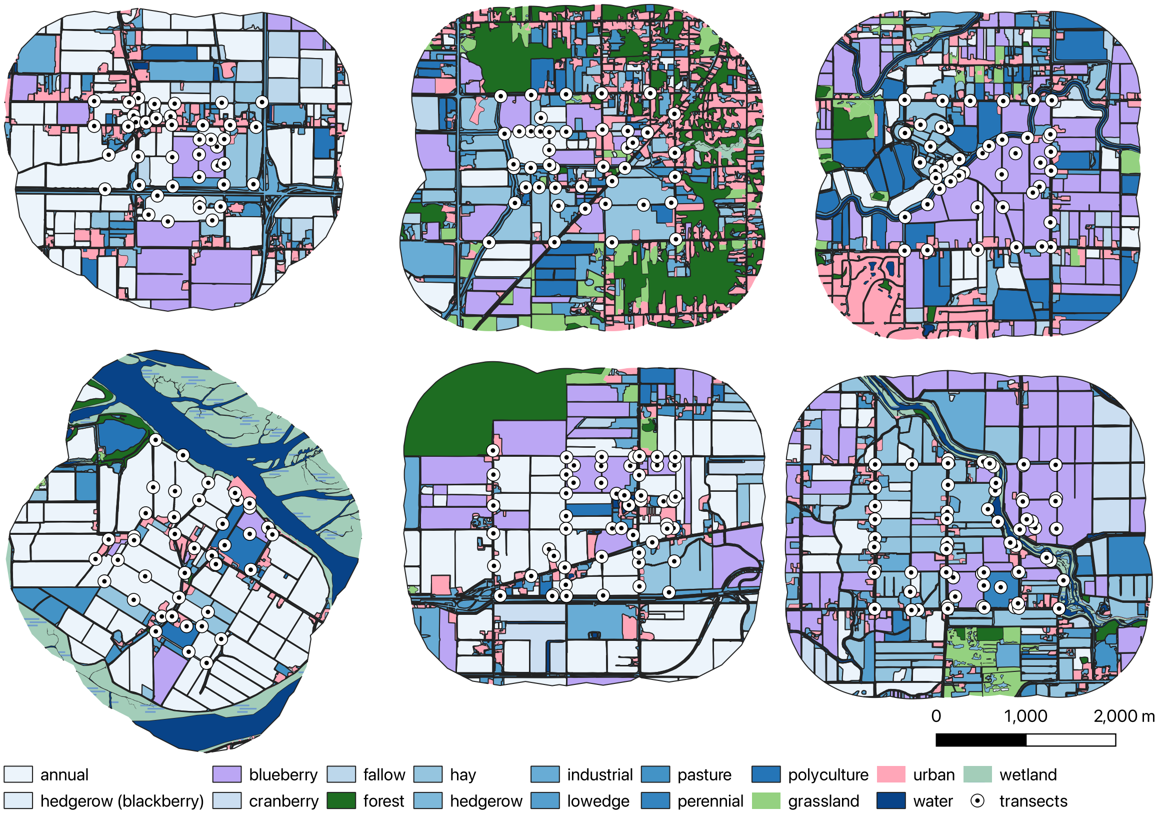


Figure A2: The six replicate landscapes, with 1000m meter buffer zones around each transect. Land cover classifications have been simplified for visibility. Black points with white circle represent the locations of transects, which were sampled for bumble bees across multiple survey rounds/years.

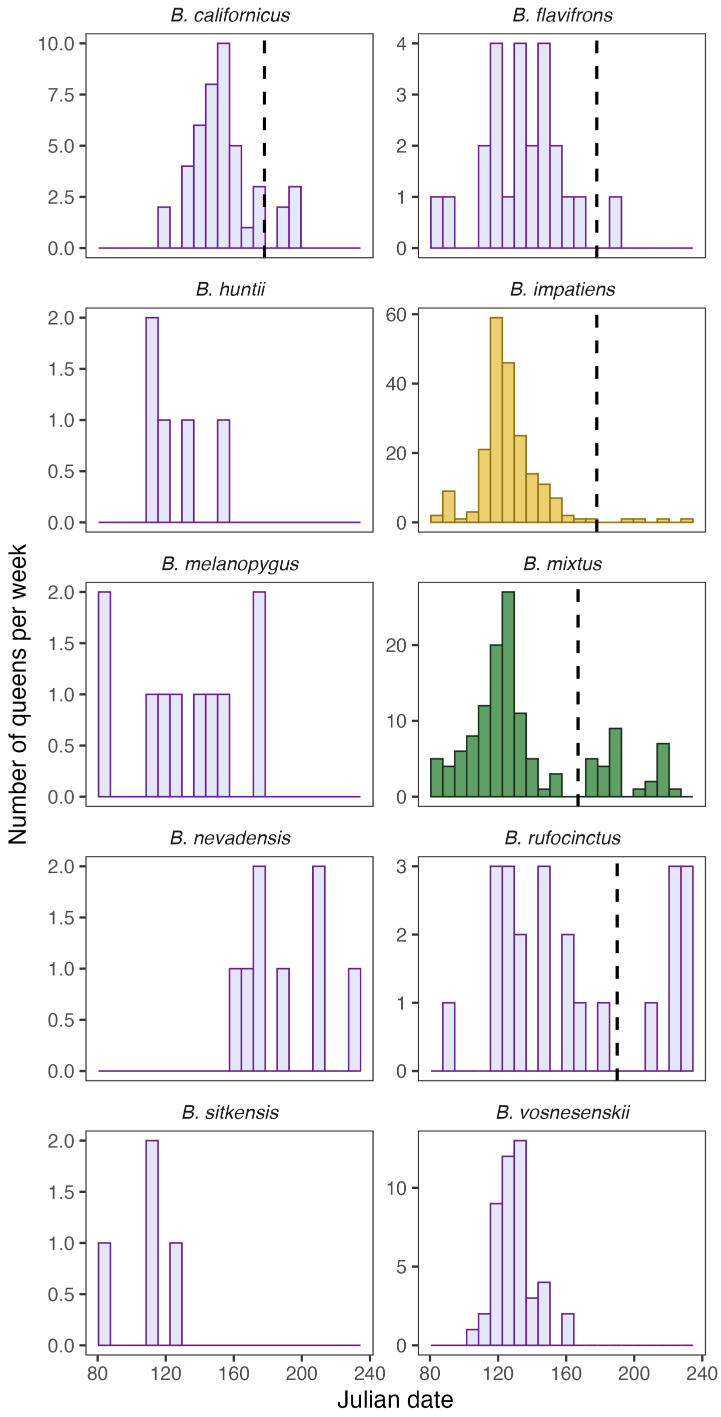


Figure A3: The number of *Bombus* queens observed each week across the sampling period. We set cut-off dates for spring queens versus summer gynes for species which showed evidence of bimodality (first mode: spring queens, second mode: summer gynes). These dates were: 15 June (*B. mixtus*), 26 June (*B. impatiens*, *B. californicus*, *B. flavifrons*), 8 July (*B. rufocinctus*).

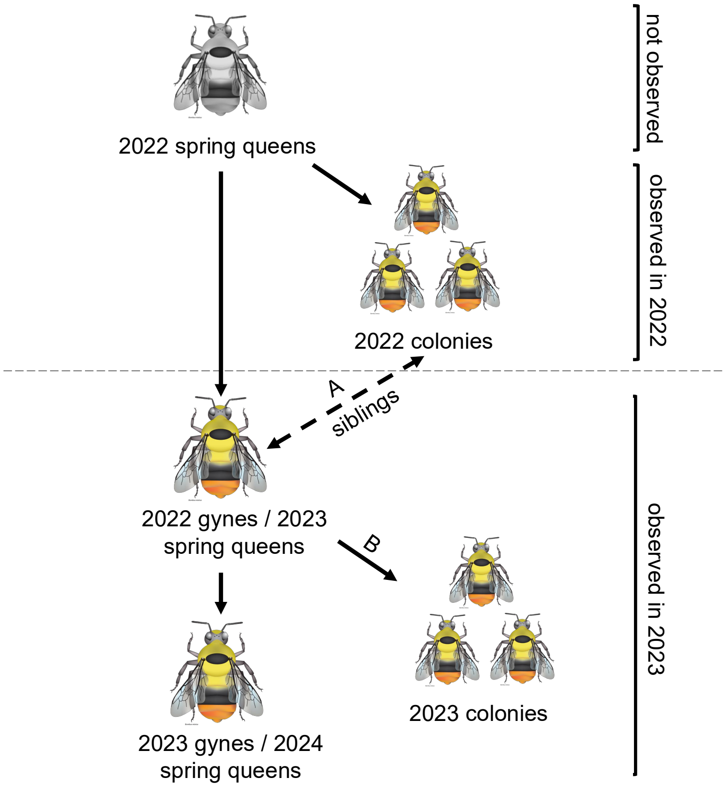


Figure A4: Family tree describing the life-stages/cohorts observed during our study. Foraging distance and colony/worker abundance models were based on observations of 2022/2023 colonymates; queen abundance and behaviour models were based on observations of 2023 spring queens. When investigating lineage turnover, we identified siblingships between 2022 colonies and 2023 spring queens (arrow A). We also searched for mother-daughter relationships between 2023 spring queens and 2023 colonies (arrow B). *Bombus mixtus* illustration from Elaine Evans and the Xerces Society, with permission.

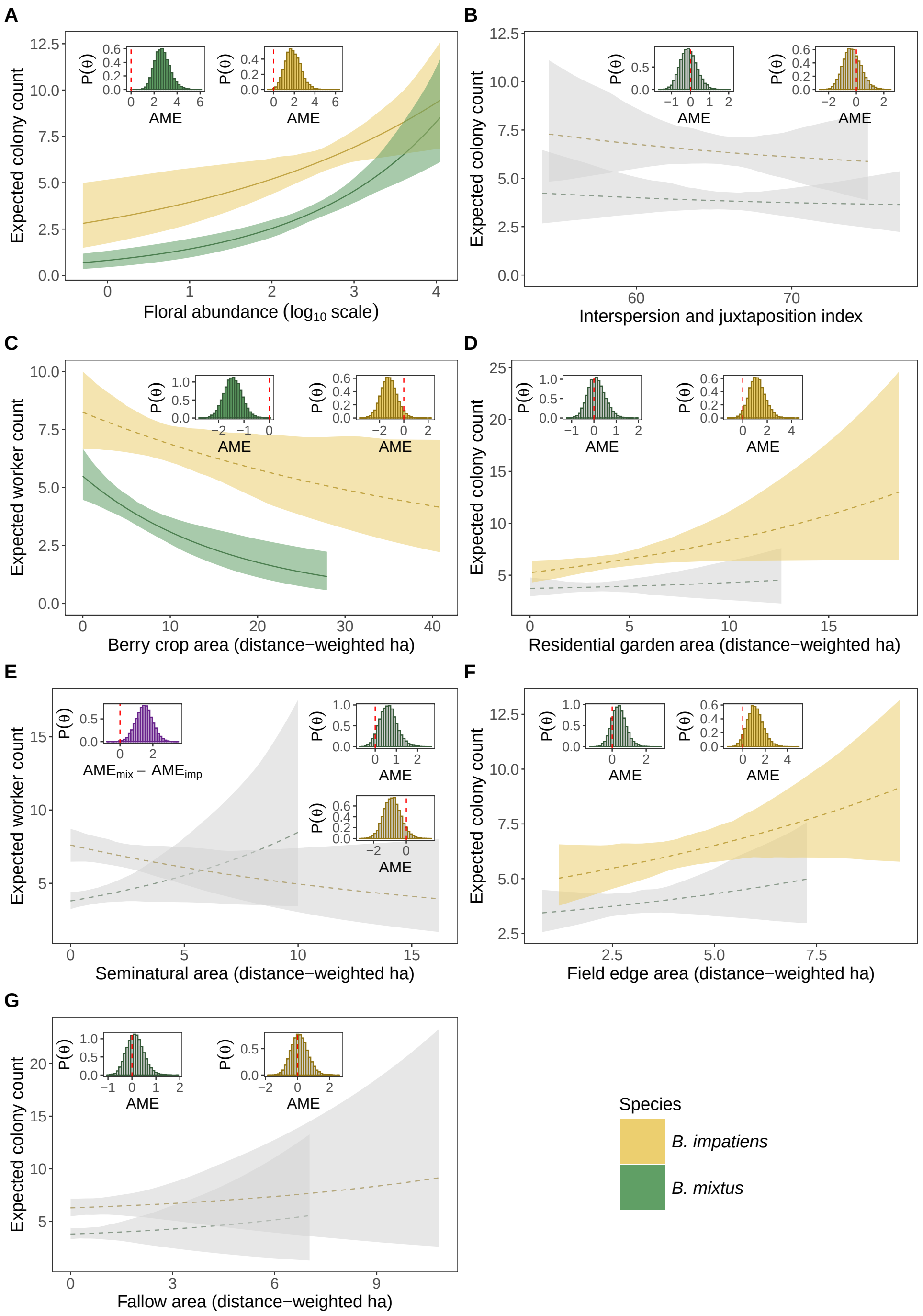


Figure A5: Worker abundance of *B. mixtus* (green) and *B. impatiens* (yellow) in relation to floral abundance (A), landscape interspersion (B), berry crop area (C), residential garden area (D), seminatural area (E), field edge area (F) and fallow arera (G). Each figure shows predicted colony counts across the range of the covariate experienced by each species. Truncated trendlines occur for *B. mixtus* because its shorter foraging range limits the maximum accessible habitat area. Shading indicates 95% credible intervals: a solid trendline and coloured shading indicated that the 95% credible interval for an association excluded zero ("strong support"); a dashed trendline and coloured shading indicates that the 90% credible interval for an association excluded zero ("moderate support"). Grey shading indicates an assocation with weak or no support. Inset figures show posteriors for the average marginal effect (AME) of each predictor. AMEs are computed for a one standard deviation increase in each predictor and are based on predictions for one hour of survey time. Vertical red dashed lines in the inset indicate zero. $P(\Theta)$: posterior density.

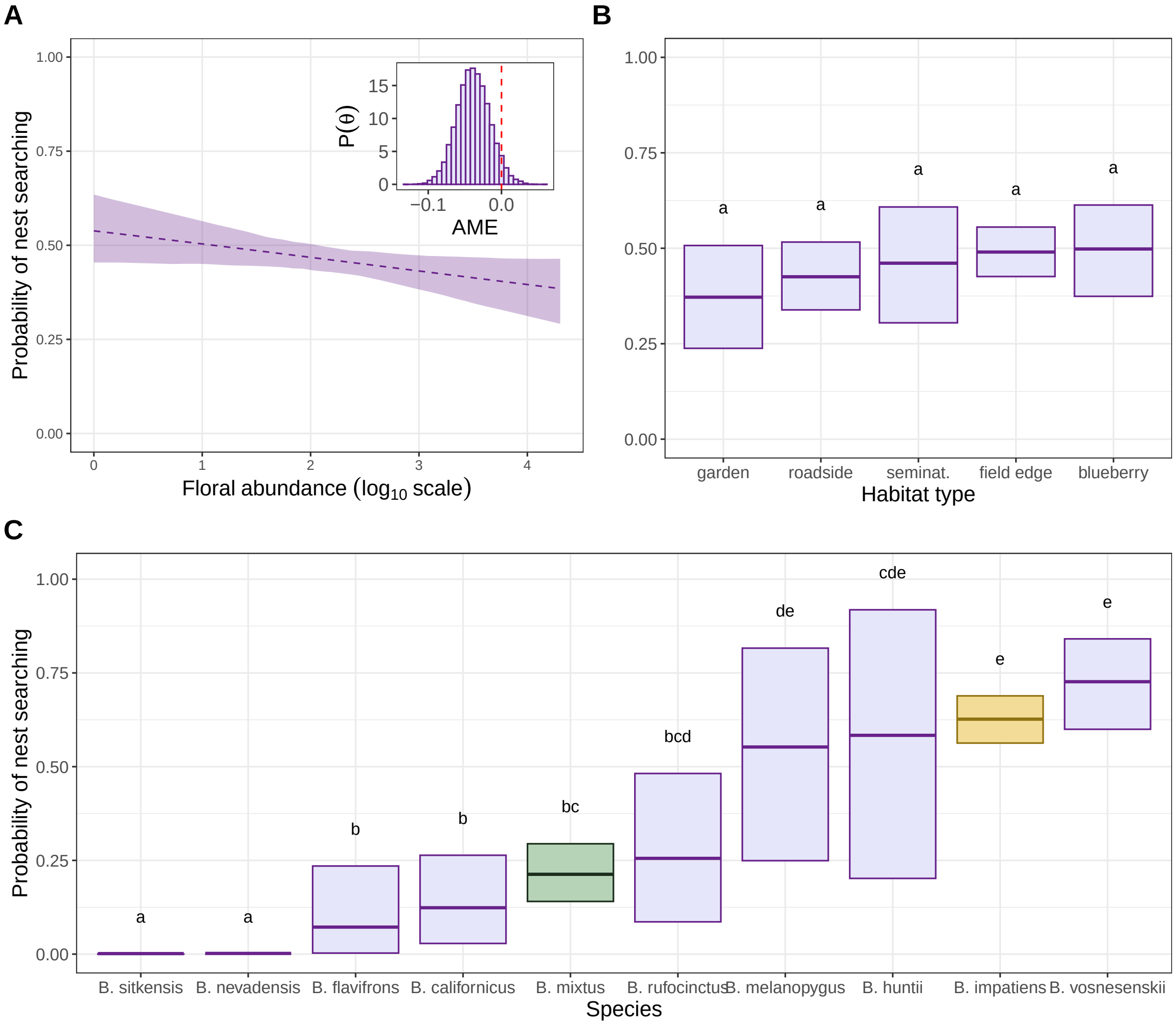


Figure A6: *Bombus* queen behaviour (1 = nest searching, 0 = foraging) in relation to (A) floral abundance, (B) habitat type, and (C) species. Shading indicates 95% credible intervals: a solid trendline and coloured shading indicated that the 95% credible interval for an association excluded zero ("strong support"); a dashed trendline and coloured shading indicates that the 90% credible interval for an association excluded zero ("moderate support"). Letters in (B-C) indicate the presence or absence of strong support for differences in the probability of nest searching between categorical levels. Inset figures show posteriors for the average marginal effect (AME) of a one standard deviation increase in $log_{10}$ floral abundance.

### A2 Population Genetics & Colony Assignments

#### A2.1 Genotyping protocol and marker information

Specimens were microsatellite genotyped at loci BT10, BTERN01, BL13, BL15, B126, BTMS0057, BTMS0059, BTMS0062, BTMS0083 (both species), BTMS0066, BTMS0086, BTMS0104, BTMS0126, BTMS0136 (*B. mixtus* only), BT28, BT30, B10, B96, B124, BTMS0073, and BTMS0081 (*B. impatiens* only) (Estoup et al., 1995, 1996; Reber Funk et al., 2006; Stolle et al., 2009). One primer for each locus was individually dye-labelled using 6FAM, NED, PET, or VIC. The majority of reverse primers were modified with 5′ PIG-tails (GTT or GTTT) to reduce stuttering. We added a 5′ cytosine (C) residue to 6FAM-labelled primers beginning with a 5′ guanidine (G) residue (e.g., BTM0126, B96, B124) to reduce 6FAM quenching. Loci were amplified in two multiplex reactions per individual. Each multiplex reaction contained 4$\mu$L of template DNA, 1$\mu$L of 10X primer mix, and 5$\mu$L of Qiagen 2X Multiplex PCR Master Mix (Qiagen, Hilden, Germany). Thermocycling was carried out following the manufacturer's protocol for Qiagen Multiplex PCR Master Mix with the following cycling conditions: initial denaturation at 95°C for 5 minutes; 35 cycles of denaturation at 95°C for 30 seconds, annealing at 57°C for 90 seconds, and extension at 72°C for 30 seconds; final extension at 68°C for 30 minutes; hold at 4°C. Diluted PCR products were submitted for automated fragment sizing at the UBC Sequencing and Bioinformatics Consortium (*B. mixtus*) and the Utah State Center for Integrated Biosystems (*B. impatiens*). 1$\mu$L of each sample was added to 8.85$\mu$L of Hi-Di formamide and 0.15$\mu$L of LIZ500 size standard; *B. mixtus* fragments were analyzed on an Applied Biosystems 3730XL 96-capillary DNA Analyzer and *B. impatiens* fragments were analyzed on an Applied Biosystems 3730 DNA Analyzer (Applied Biosystems, Foster City, CA, USA). Microsatellite peaks were assigned manually using the software Geneious Prime 2024.0.5 (https://www.geneious.com). Scoring error rates were assessed by re-genotyping a panel of 96 individuals per species, and loci with observed error rates ≥3% were discarded from further analyses (BL15 for both species).

Rarified allelic richness was calculated for a single pooled population using rarefaction to the locus-specific number of observed gene copies (i.e., accounting for missing genotypes) following Hurlbert's formulation as implemented in *hierfstat* (Goudet, 2005). The number of individuals genotyped per population was 3588 for *B. mixtus* and 5012 for *B. impatiens*.

See Table A9 for primer sequences, size ranges, rarified allelic richness (RAR), multiplexing information, and locus retention status following quality screening.

Table A9: Description of loci used for microsatellite genotyping of *B. mixtus* and *B. impatiens*. **RAR** = rarified allelic richness, **GTTT** = addition of PIG-tail.

| Locus | Forward Sequence | Reverse Sequence | Species | Size Range | RAR | Plex | Dye Tag | Maintained |
|---|---|---|---|---|---|---|---|---|
| **BTMS0086** | CTCGCGCTTGTCGAATCAAT | **GTTT**AGAGAAATTGCATGCGGTCG | *B. mixtus* | 266-277 | 3 | 1 | VIC | yes |
| | | | *B. impatiens* | - | - | - | - | - |
| **BTMS0057** | TGCTTGAACCGAAATAGAGGG | **GTTT**CACCGGCATTTTACACACCA | *B. mixtus* | 116-136 | 11 | 1 | VIC | yes |
| | | | *B. impatiens* | 106-136 | 14.6 | 2 | VIC | yes |
| **BTMS0136** | GCATTCGGGTATTGCGTTCTTTAG | **GTTT**CGTTTATCTGCTTCTCTCGTTCG | *B. mixtus* | 154-196 | 20 | 1 | PET | yes |
| | | | *B. impatiens* | - | - | - | - | - |
| **BTMS0066** | TTAACGCCCAATGCCTTTCC | CATGATGACACCACCCAACG | *B. mixtus* | 113-197 | 26.9 | 1 | NED | yes |
| | | | *B. impatiens* | - | - | - | - | - |
| **BTMS0062** | CTGGGCGTGATTCGATGAAC | **GTTT**CTGTCGCATTATTCGCGGTT | *B. mixtus* | 229-289 | 28 | 1 | NED | yes |
| | | | *B. impatiens* | 230-330 | 34.9 | 1 | NED | yes |
| **BT10** | TCTTGCTATCCACCACCCGC | **GTTT**GGACAGAAGCATAGACGCACCG | *B. mixtus* | 128-172 | 22 | 1 | 6FAM | yes |
| | | | *B. impatiens* | 119-157 | 27.8 | 1 | 6FAM | yes |
| **BTMS0104** | TCCTCTGTTCAGCACACGAT | **GTTT**TTCGAAGCCTCGATGTCGT | *B. mixtus* | 266-268 | 2 | 1 | 6FAM | no ($F_{is}$) |
| | | | *B. impatiens* | - | - | - | - | - |
| **BL15** | CGAACGAAAACGAAAAAGAGC | TCTTCTGCTCCTTTCTCCATTC | *B. mixtus* | 124-172 | - | 2 | VIC | no (error rate) |
| | | | *B. impatiens* | 137-173 | - | 1 | VIC | no (error rate) |
| **B126** | CGATTCTCTCGTGTACTCC | **GTTT**GCTTGCTGGTGAATTGTGC | *B. mixtus* | 143-151 | 7 | 2 | PET | yes |
| | | | *B. impatiens* | 142-200 | 17.5 | 1 | PET | yes |
| **BL13** | CGAATGTTGGGATTTTCGTG | **GTTT**GCGAGTACGTGTACGTGTTCTATG | *B. mixtus* | 158-214 | 23 | 2 | NED | yes |
| | | | *B. impatiens* | 148-184 | 8.9 | 1 | NED | no (LD) |
| **BTMS0083** | CGACTCGTTCGAGCGAAATTA | GTTTTTGCCAGGCTCCGAAT | *B. mixtus* | 266-304 | 20 | 2 | NED | yes |
| | | | *B. impatiens* | 276-320 | 21 | 2 | NED | yes |
| **BTERN01** | CGTGTTTAGGGTACTGGTGGTC | **GTTT**GGAGCAAGAGGGCTAGACAAAAG | *B. mixtus* | 104-120 | 10 | 2 | 6FAM | no ($F_{is}$) |
| | | | *B. impatiens* | 101-151 | 15.9 | 2 | 6FAM | yes |
| **BTMS0059** | AGTTCGACAGACCAAGCTGT | **GTTT**GGCTAGGAAAGATTAGCACTACC | *B. mixtus* | 342-362 | 6 | 2 | 6FAM | no ($F_{is}$) |
| | | | *B. impatiens* | 339-367 | 8.9 | 1 | 6FAM | yes |
| **BTMS0126** | **C**CGGTGATCGCTTAAAGCTC | **GTTT**GCCAACTACGTTCAATATCG | *B. mixtus* | 163-195 | 18 | 2 | 6FAM | yes |
| | | | *B. impatiens* | - | - | - | - | - |
| **B96** | **C**CGGGAGAGAAAGACCAAG | **GTTT**GATCGTAATGACTCGATATG | *B. mixtus* | - | - | - | - | - |
| | | | *B. impatiens* | 236-278 | 18 | 1 | PET | yes |
| **BTMS0081** | ACGCGCGCCTTCTACTATC | **GTT**AGGGACACGCGAACAGAC | *B. mixtus* | - | - | - | - | - |
| | | | *B. impatiens* | 292-320 | 6.8 | 1 | VIC | yes |
| **B10** | GTGTAACTTTCTCTCGACAG | **GTTT**GGGAGATGGATATAGATGAG | *B. mixtus* | - | - | - | - | - |
| | | | *B. impatiens* | 184-256 | 22.5 | 2 | NED | yes |
| **BT28** | TTGCTGACGTTGCTGTGACTGAGG | **GTT**TCCTCTGTGTGTTCTCTTACTTGGC | *B. mixtus* | - | - | - | - | - |
| | | | *B. impatiens* | 177-207 | 10.7 | 2 | PET | no ($F_{is}$) |
| **BT30** | ATCGTATTATTGCCACCAACCG | **GTT**CAGCAACAGTCACAACAAACGC | *B. mixtus* | - | - | - | - | - |
| | | | *B. impatiens* | 173-206 | 9.8 | 2 | VIC | yes |
| **B124** | **C**CGCAACAGGTCGGGTTAGAG | **GTTT**CAGGATAGGGTAGGTAAGCAG | *B. mixtus* | - | - | - | - | - |
| | | | *B. impatiens* | 233-303 | 19.8 | 2 | 6FAM | yes |
| **BTMS0073** | CGATATCGCGATCTTCGTACAC | **GTT**GTAGCATGCTCTCCGTGTTG | *B. mixtus* | - | - | - | - | - |
| | | | *B. impatiens* | 112-136 | 7.9 | 2 | PET | no ($F_{is}$) |

#### A2.2 Assessing locus $F_{is}$, $F_{st}$ and linkage disequilibrium

We used an iterative approach to assign workers and queens to their natal colonies. First, using the pedigree reconstruction software COLONY 2.0.6.5 (Jones & Wang, 2010) (hereafter, COLONY), we assigned full siblingships based on all available microsatellite data, assuming male and female monogamy and no inbreeding. A single run was carried out for each species and year, using the software's full-likelihood approach and no siblingship size scaling or priors. Sibling pairs were maintained when $P_{fullsibdyad} = 1$. A single individual from each putative colony (including non-circular colonies, described below) was maintained for downstream analyses of locus quality.

While COLONY can account for inbreeding at the population level, locus-specific estimates of $F_{is}$ should generally be similar to one another for a given population, reflecting their shared evolutionary history. Loci with $F_{is}$ estimates that deviate significantly from the species mean (across loci) may suffer from null alleles or other types of scoring errors that can bias siblingship assignment. We therefore tested individual locus deviations from population mean $F_{is}$ as a criterion for marker inclusion/exclusion. Single locus $F_{is}$ estimates were computed for all landscape, year, species groups following Weir and Cockerham (1984) in the package *genepop* (Rousset, 2008) (Fig. A7). We then fit linear models to $F_{is}$ estimates with locus and landscape as fixed predictors. We used sum-to-zero coding so that model intercepts represented mean $F_{is}$ across all loci, and calculated the estimated marginal mean of each locus using the *emmeans* package (Lenth, 2024) (Fig. A8A-B). We iteratively removed loci with $F_{is}$ significantly different from the species mean $F_{is}$, starting with the locus with the greatest deviation, and re-running the model after each removal. We did not apply an adjustment for multiple-hypothesis testing, but instead utilized a relatively stringent p-value ($\alpha$ = 0.01) for removal of loci. This process resulted in the removal of loci BTERN01,

BTMS0104, and BTMS0059 from downstream analyses for *B. mixtus* and removal of BTMS0073 and BT28 for *B. impatiens*. $F_{is}$ estimates for the remaining loci (n = 10 for *B. mixtus*, n = 13 for *B. impatiens*) can be found in Fig. A8 C-D.

Next, we checked locus pairs for linkage disequilibrium (LD). Marker linkage can lead to non-independent assortment, a condition which violates the assumptions of most parentage reconstruction software and can result in overconfidence in estimated sibling pairs. LD was calculated for each locus pair in each population (landscape, year, species) in *genepop*. We applied a Bonferroni correction for multiple hypothesis testing, and flagged locus pairs which showed significant deviations in $> 2$ populations. For *B. mixtus*, four locus pairs showed signs of LD, but each occurred at only a single landscape in a single year, so we chose to maintain all loci. For *B. impatiens*, there was strong evidence for linkage disequilibrium between BTMS0057 and BL13 (7 out of 12 populations showed significant LD following Bonferroni correction). To determine which locus to maintain, we calculated the polymorphic information content (PIC) using the package *PopGenUtils* (Tourvas, 2025). We found that BTMS0057 had higher PIC in both years (PIC = 0.78) compared to BL13 (PIC = 0.47 in 2022 and 0.45 in 2023). We therefore maintained BTMS0057 and removed BL13 from further analyses for *B. impatiens*.

After locus quality screening, we calculated global and pairwise $F_{st}$ following Nei (1987) in the package *hierfstat* (Goudet, 2005). For both species and years, estimates of global $F_{st}$ were less than 0.005. Pairwise $F_{st}$ ranged from -0.002 to 0.01, indicating little or no genetic differentiation between surveyed sub-populations. Finally, we computed the marginal mean $F_{is}$ for each landscape to determine whether there was evidence for inbreeding following locus removal, and whether it varied between landscapes (Fig. A9). Because we found evidence for only minor inbreeding (e.g., $F_{is} < 0.05$) we ran all final colony assignments using the no-inbreeding model. Given the very low estimates of global and pairwise $F_{st}$, we chose to combine landscapes for siblingship assignments to maximize the accuracy of allele frequency estimation.

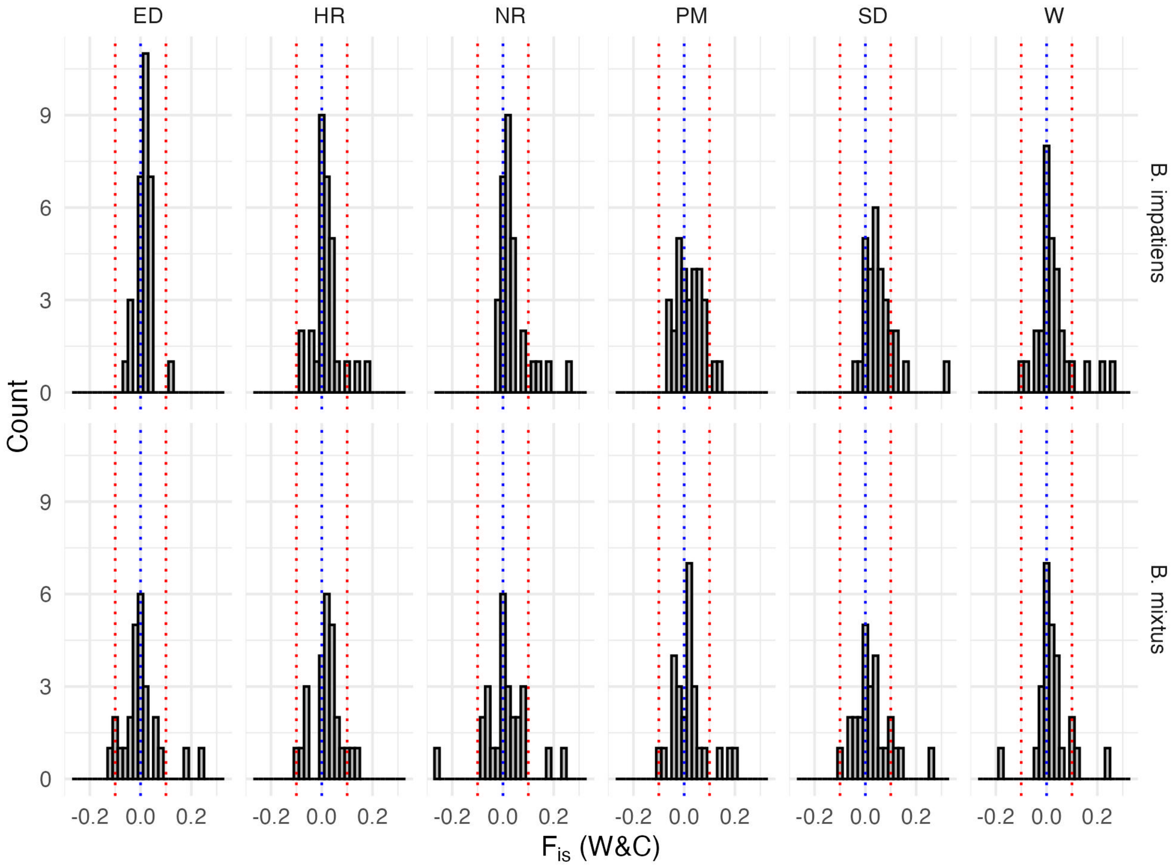


Figure A7: Estimates of $F_{is}$ for each locus in each subpopulation. Estimates from 2022 and 2023 were calculated separately but are shown together for each site x species combination. Blue dotted lines indicates $F_{is} = 0$ and red dotted lines indicate $F_{is} = \pm 0.1$.

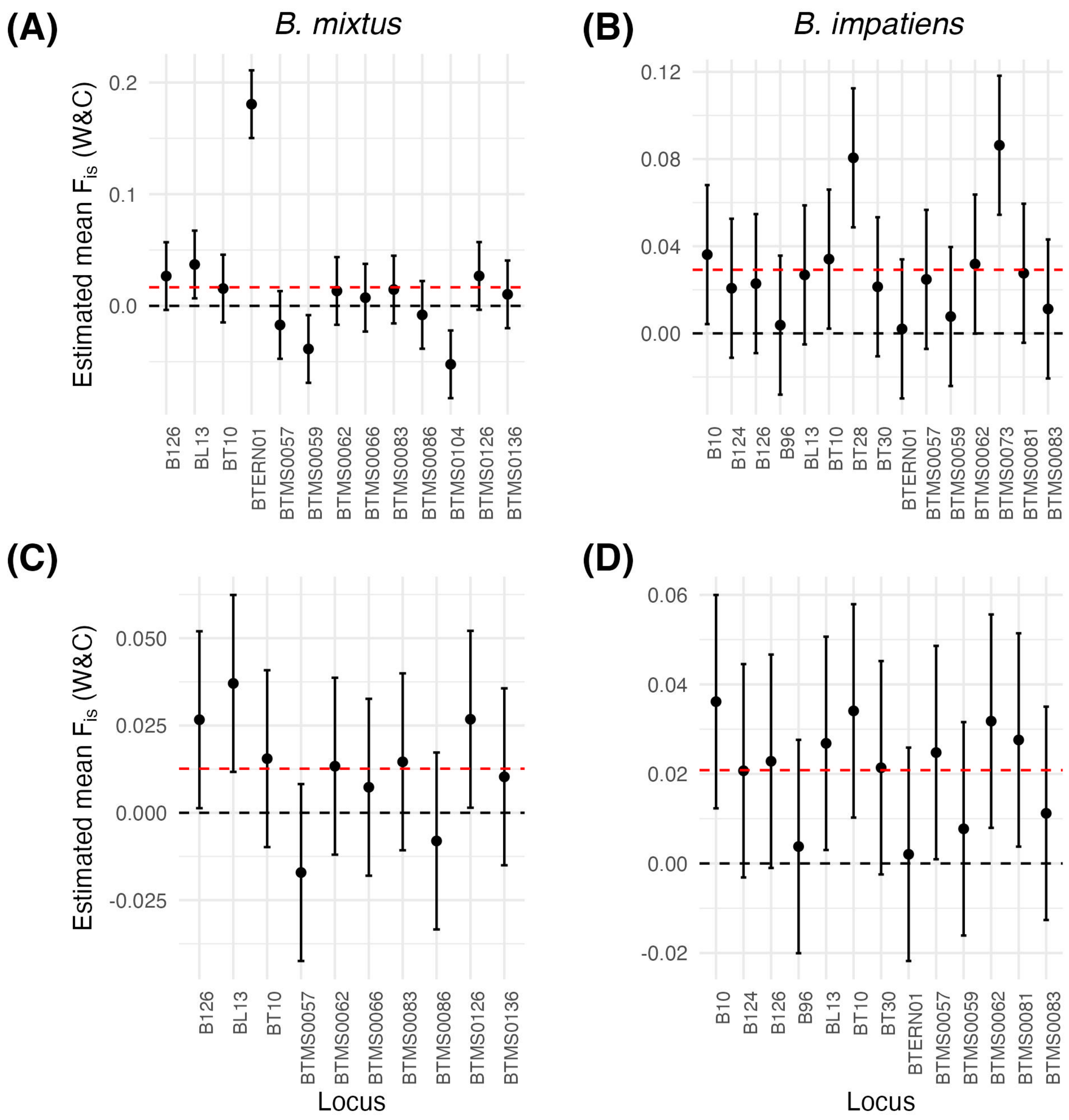


Figure A8: Locus-specific $F_{is}$ marginal means. A) *B. mixtus* all loci; B) *B. impatiens* all loci; C) *B. mixtus* loci following iterative removal of loci which differed significantly from global mean $F_{is}$; D) *B. impatiens* loci following iterative removal of loci which differed significantly from global mean $F_{is}$. Dashed black line denotes $F_{is} = 0$, dashed red line denotes global mean $F_{is}$ for each species.

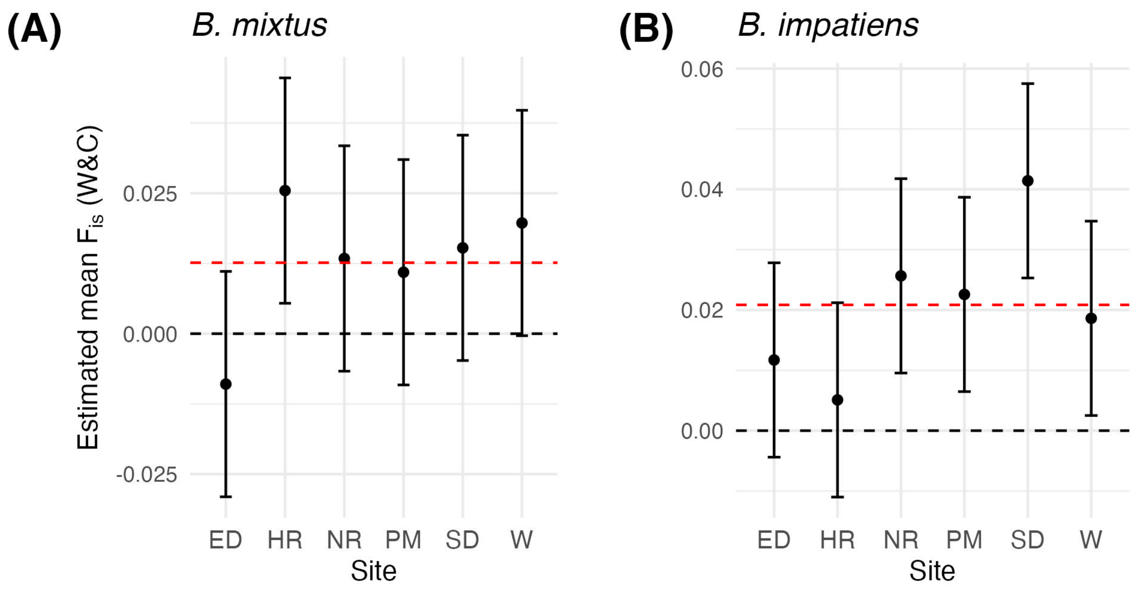


Figure A9: Landscape-specific $F_{is}$ marginal means following removal of low-quality loci for A) *B. mixtus* and B) *B. impatiens*. Dashed black line denotes $F_{is} = 0$, dashed red line denotes global mean $F_{is}$ for each species.

#### A2.3 Testing COLONY on simulated data

To test the informativeness of our genetic loci and validate the accuracy of COLONY 2.0.6.5 (Jones & Wang, 2010) for detecting siblingships amongst our specimens, we performed colony assignments on multiple simulated datasets using realistic family sizes, spatial distributions, and allelic frequencies.

We approached these simulations with four objectives:

(i) To determine false positive and false negative siblingship assignment rates, given the informativeness of our microsatellite datasets,

(ii) To inform an appropriate strategy (probability threshold, number of runs of the software) for maintaining or rejecting each sibling pair;

(iii) To select suitable software parameters, and in particular to evaluate the usefulness of siblingship size priors and exclusion of between-landscape siblingships for reducing false positive rates;

(iv) To assess whether our dataset(s) are sufficiently informative to support identification of female polygamy, which has been observed in North American *Pyrobombus* (Owen & Whidden, 2013; Payne et al., 2003).

#### A2.3.1 Simulation strategy

**Spatially explicit siblingships** We first simulated spatially explicit siblingships following Pope and Jha (2017).

We began by simulating six 5 x 5 trapping grids (traps $k \in \kappa$) on a single raster surface comprised of cells $j \in \mathbb{J}$. Colonies $i \in \mathbb{C}$ were distributed uniformly at random throughout the "landscape" (Fig. A10 A-B).

We then sampled individuals from colonies $i \in \mathbb{C}$ captured at traps $k \in \mathbb{K}$ from the joint distribution $\Pr(s, c \mid s \in \kappa)$, where $\{s, c\}$ are the indices of a random visitation event of an individual from colony $c \in \mathbb{C}$ to grid cell $s \in \mathbb{J}$. Details of the sampling process can be found in (Pope & Jha, 2017).

We define the foraging kernel of workers from colony $i$ as

$$\Pr(s = k \mid c = i) \ = \ \frac{\lambda_i(k)}{\sum_{j \in J} \lambda_i(j)} \tag{4}$$

where the visitation intensity of individuals from colony $i$ to location $j$ is

$$log(\lambda_i(j)) = \frac{-\|x_j - \delta_i\|}{\rho} \tag{5}$$

$x_j$ are the spatial coordinates of any grid cell in the raster, and $\delta_i$ are the spatial coordinates of colony $i$. The foraging kernel in this example is therefore symmetrical and exponentially decaying as a function of distance from the colony location.

For each simulation, samples are drawn until a stopping point (desired number of samples) is reached. $n_i$ (the number of individual per colony) is updated after each "sampling event" to prevent oversampling from colonies located very close to traps.

To verify that the size of sampled siblingships (e.g., number of siblings per sibling group) accurately mirrors the distribution of siblingship sizes in real data, we compared our simulated distributions to the distribution of siblingshp sizes in our real data (Fig. A10 C-D). We found that moderating the background density of colonies (i.e., the total number of colonies simulated on the landscape) was the most effective strategy for controlling average siblingship size. A higher background density of simulated colonies results in a higher proportion of singleton colonies (colonies represented by only a single individual).

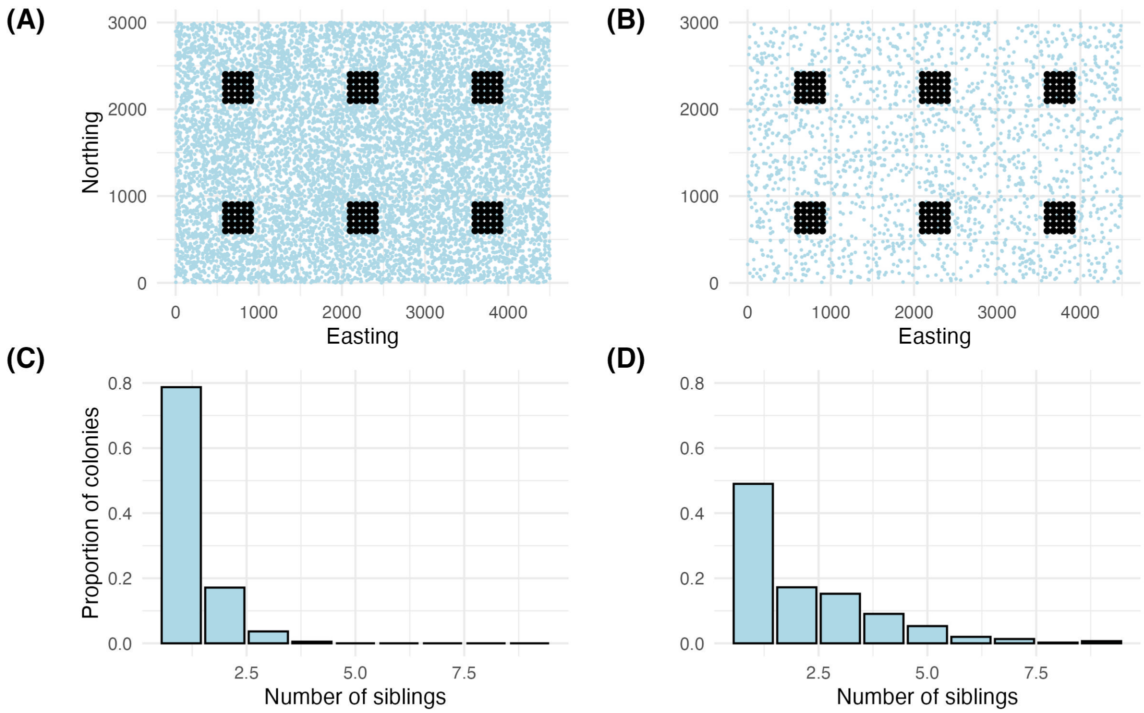


Figure A10: Simulations of spatially explicit siblingships. (A-B) Spatial distribution of traps (black points) and simulated colonies (light blue points). (C-D) Distribution of siblingship sizes in final simulated datasets. (A,C) Simulations using 10,000 background colonies. (B,D) Simulations using 2,000 background colonies. In general, a higher background density of colonies results in a smaller average family size; in both cases, some colonies are not observed in the final dataset. Each map unit in our simulation represents 5 meters (e.g., 1000 map units = 5000 meters). The background colony densities represented here therefore equate to 0.26 colonies per hectare and 0.06 colonies per hectare; we use the former value, as it is in line with previous estimates of landscape-wide colony densities and creates a more realistic distribution of siblingship sizes (C).

**Multilocus genotypes** We simulated multilocus genotypes for each sampled individual under several mating scenarios. In the simplest case, we assume monogamy for both males and queens. The majority of the simulation results presented below follow this assumption. In a second set of simulations, we assumed varying rates of polyandry (e.g., queen polygamy) to assess the impact of this paradigm on siblingship inference. For each simulation we used the following heuristic:

1. Simulate parental genotypes for each siblingship based on the allele frequencies present in our real data. These frequencies were inferred from an earlier run of COLONY, which provides estimated frequencies after accounting for height-

ened frequency of alleles present in large families; because average family size was small in our dataset (< 2 individuals) and large families were rare (Fig A10 C), raw allele frequencies would have likely been sufficient.

2. Randomly draw offspring genotypes from the set of possible parental alleles at each locus. For simulations involving monogamous mating, father haplotypes are assigned directly to all offspring in the siblingship; in colonies with multiple paternity, we assume two fathers and assign inheritance of paternal haplotypes from $\Pr(father_1, father_2) = (0.7, 0.3)$ following the proportions observed for *B. impatiens* in Bird et al. (2024).

3. Introduce errors and data missingness based on observed rates for our real datasets. To introduce errors, we mutate each allele with a probability equal to the rate of errors for that locus and species; we assume that most errors are due to contamination, rather than allele dropout, and therefore draw new (erroneous) alleles from the allele frequency distribution for each species. We observed that individuals which were missing data for *one* copy of a locus were more likely to be missing data for *both* copies than if missingness were distributed uniformly at random. This is likely because there were two primary missingness-generating processes in real data: amplification failure (both alleles missing for an individual) and binning failure (one or both alleles missing for an individual). (In cases where only one copy of a locus failed to amplify, heterozygous individuals would be falsely classified as homozygous—an error, rather than missing data). To mimic the observed distribution of missingness, we first calculate the proportion of missing data for each marker ($P_{missing}$) and then remove data for (1) both alleles, with probability $1/3 * P_{missing}$ and for (2) a single allele, with probability $1/3 * P_{missing}$.

This method allows us to draw conclusions based on the informativeness of our specific genetic datasets, rather than an idealized situation with perfect data. We performed simulations based on allele frequencies for both species (*B. mixtus* and *B. impatiens*) because variation in marker number and/or polymorphic information content could lead to differing results.

##### A2.3.2 Determining an appropriate heuristic for maintaining or rejecting inferred siblingships

Like any software for family reconstruction, COLONY can produce erroneous siblingships (false positives) or fail to identify kinship when it exists (false negatives). Our preliminary analyses resulted in a high number of inferred siblingships between individuals separated by >20 km when individuals from all study landscapes were permitted to form siblingships. While the biology of bumblebee foraging/dispersal does not unilaterally exclude the possibility of such distant relationships, the likelihood of observing such separation distances is extremely small, and unlikely to represent biological reality except in very rare cases.

A common strategy in studies performing *Bombus* colony assignment is to repeat multiple "runs" (usually 2-5) of the COLONY software on the dataset, and maintain family groups which are inferred in all runs at or above some confidence threshhold (usually $P \geq 0.95$, but sometimes $P \geq 0.8$). See, for example, Carvell et al. (2012, 2017), Dreier et al. (2014), Mola et al. (2020), and Rao and Strange (2012). However, we are not aware of any studies which give support for a particular threshhold probability or number of runs necessary to reach a particular confidence level in assignments, nor to achieve a satisfactory balance between false positive siblingships and false negative siblingships. Indeed, the desirable threshhold is likely to vary as a function of the number and informativeness of markers for a given population.

To overcome these limitations, we tested probability exclusion criteria from $P = 0.95$ to $P = 1$, for 1 or 5 runs of COLONY version 2.0.6.5. Further, we compared the use of family cluster probabilities (COLONY output file .BestCluster—hereafter referred to as the family method) and full sibling dyad probabilities (COLONY output .FullSibDyad—hereafter referred to as the dyad method).

We began by simulating 5 datasets (e.g., different siblingship arrangements with unique parental genotypes) consisting of n = 1200 individuals each, which was roughly the midpoint of population sizes for our real data. For each dataset we performed 5 runs of COLONY (see Table A10 for a summary of COLONY software settings).

Based on our results, we conclude that the software converges reliably for datasets like ours, and that repeated runs of the software have little or no effect on conclusions drawn. In most cases, false positive and false negative rates are either identical or nearly overlapping, regardless of the numbers of runs (Fig A11, A12). We therefore recommend that to save time and computational resources, researchers should check for convergence for each microsatellite dataset using 2-3 runs of the software, and if convergence is achieved they should feel confident that a single run is sufficient to identify siblingships.

For both species we found that increasing the probability threshhold from 0.95 to 1 more effectively reduced false positives for the dyad method than for the family method, although in general a probability threshold $\geq 0.99$ was necessary to maintain false positive rates at around 5% using the dyad method. For $P = 1$ the dyad method led to a high proportion of false negatives ($\geq 5\%$) (Fig A12). For this reason, we would not recommend using this maximal stringency unless a very low rate of false positives is required.

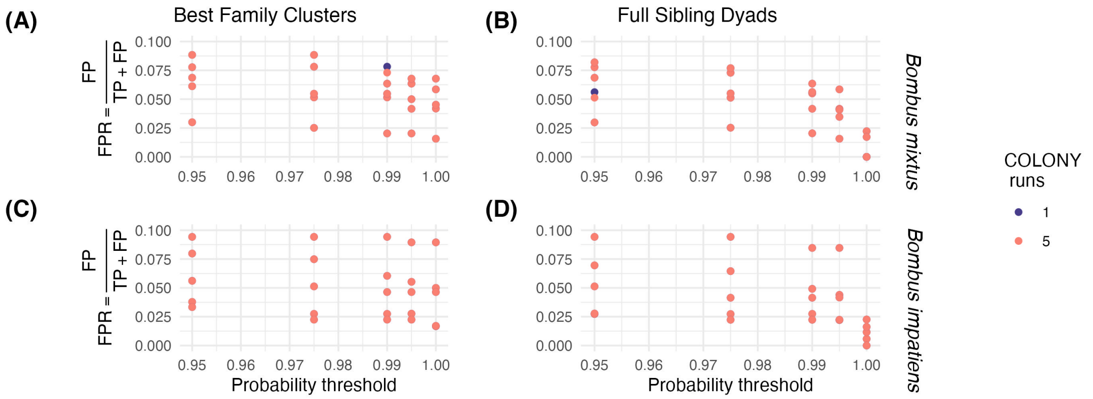


Figure A11: False positive rates produced by COLONY 2.0.6.5 when assigning siblingships to simulated data based on *B. mixtus* (A-B) and *B. impatiens* (C-D) allele frequencies and marker numbers. Shown here as a function of the probability threshold used to maintain siblingships. (A, C): False positive rates when siblingships are assigned via the “family" method (using .BestCluster output), (B, D): False positive rates when siblingships are assigned via the “dyad" method (using .FullSibDyad output). Color denotes the number of replicated runs of the software used to establish siblingships; in most cases, points are perfectly overlapping.

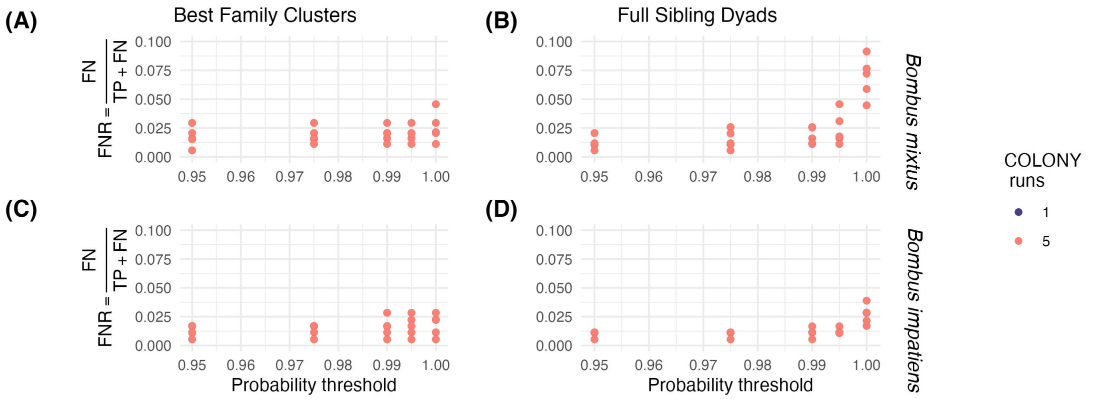


Figure A12: False negative rates produced by COLONY 2.0.6.5 when assigning siblingships to simulated data based on *B. mixtus* (A-B) and *B. impatiens* (C-D) allele frequencies and marker numbers. Shown here as a function of the probability threshold used to maintain siblingships. (A, C): False negative rates when siblingships are assigned via the “family" method (using .BestCluster output), (B, D): False negative rates when siblingships are assigned via the “dyad" method (using .FullSibDyad output). Color denotes the number of replicated runs of the software used to establish siblingships; in most cases, points are perfectly overlapping.

#### A2.3.3 Handling “non-circular" families

To fully assess utility of the family method versus the dyad method, we need a method for resolving “non-circular" families. They families arise when some (but not all) individuals in a group are inferred to be full siblings (e.g., A related to B, B related to

C, A not related to C). Such cases are rare, and handled internally by COLONY to create *family clusters* that are circular. If we make assignments based on probabilities of each dyad pair, we are left the task of deciding how to resolve these non-circular families. Indeed, resolution of non-circular families could be one process which leads to the higher false positive rates previously observed when using the family method (Fig A11).

We first explored the structure of non-circular families in our simulated datasets, to determine whether non-circularity is more frequently the result of false positives (e.g., a third individual being erroneously added to a sibling pair) or false negatives (e.g., failure to detect a sibling relationship between any pair of siblings in a triad).

To do this, we identified non-circular families from five simulations of 2000 individuals each, using a threshold of $P = 0.995$ for inclusion of pairwise relationships. When then classified each missing link as either a false negative (a true siblingship that was not inferred by our method) or a true negative (a false siblingship that was correctly excluded, meaning that at least one of the other siblingships in the non-circular family was a false positive). Fig A13 shows the distribution of false negatives and true negatives in simulated datasets based on *B. mixtus* and *B. impatiens* allele frequencies.

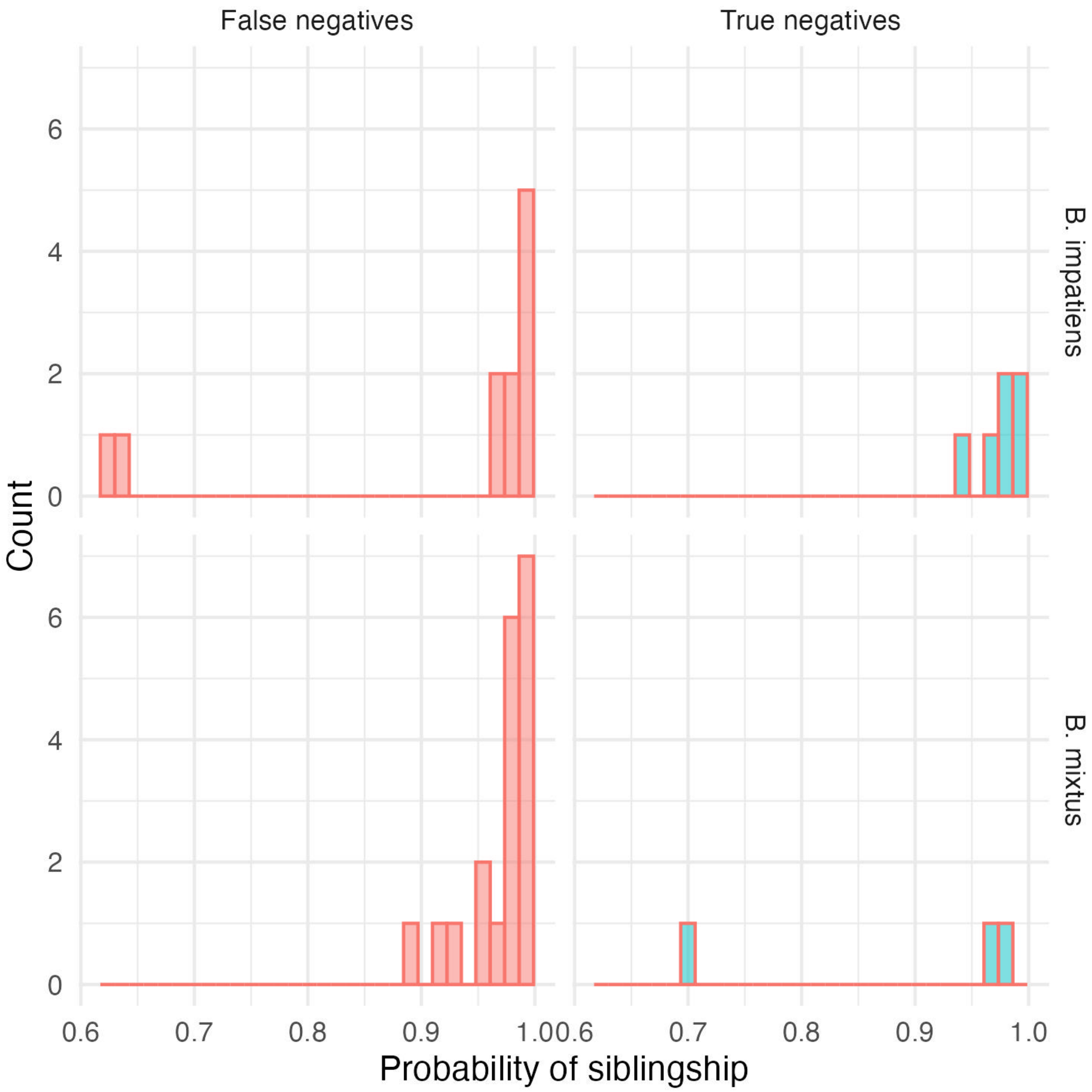


Figure A13: Frequency of false negative and true negative “missing links" in non-circular siblingships. The x-axis shows the probability assigned to each missing link by the COLONY full sib dyad method (maximum probability for missing links = 0.994, probability threshold for initial acceptance of sibling pairs = 0.995).

While there does not appear to be a convenient threshold at which we are able to exclude all true negatives while accepting all false negatives, it is comforting that the majority of missed links appear to be false negatives, meaning that we can select a lower probability threshold (e.g., $P = 0.95$) at which to accept these relationships and circularize families without introducing a large number of false positives.

After this step, we are left with a small number of remaining non-circular families (e.g., missing links rejected even at a lower probability threshold). We resolve these families by maintaining the largest “clique" (e.g., group in which all siblings are connected) or by randomly selecting one complete clique, if there are multiple of equal size.

#### A2.4 Performance of family and dyad methods with and without siblingship size priors

When comparing the family method to the dyad method for siblingship assignment, we found that results were strongly dependent on whether a siblingship size prior was used. The siblingship prior allows researchers to set a prior on the harmonic mean family size, based on their previous understanding of the expected family sizes in their dataset or similar datasets. Based on our preliminary analyses, about 70-80% of individuals in our true dataset originated from “singleton" colonies (no siblings in the dataset), so we set our siblingship priors to 1 individual per family.

For this analysis, we simulated datasets of N = 2000 individuals and subsampled 20%, 40%, 60%, 80%, or 100% of individuals to test the use of siblingship priors across datasets of multiple sizes. Our probability threshold was $P = 0.995$.

When not utilizing a prior, we found that the family method was much less reliable than the dyad method for excluding erroneous siblingships (Fig A14); false negative rates were similar for both strategies. For the family method, 10-60% of all inferred pairwise relationships were false positives when not using a size prior. False positive rates were consistently $\leq 0.2$ when a prior was used (Fig A14). The dyad method, in contrast, resulted in $\leq 12\%$ false positives for all conditions tested, and was typically $\leq 5\%$ (Fig A15). Interestingly, we found that when we assigned siblingships based on the dyads method, the use of a siblingship size prior has little effect on the false positive or false negative rates for *B. mixtus*, but caused a marginal increase in false positives and a marginal decrease in false negatives for *B. impatiens* (Fig A15).

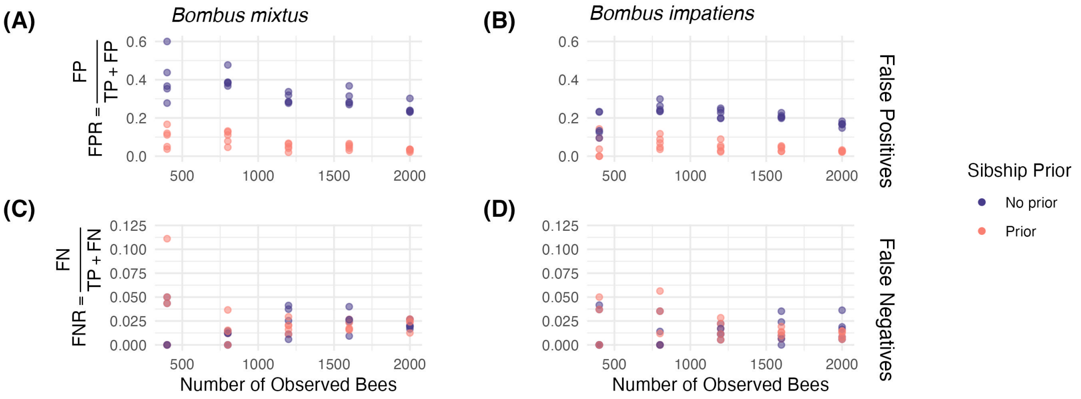


Figure A14: False positive (A-B) and false negative (C-D) rates of siblingship assignment in *B. mixtus* (A, C) and *B. impatiens* (B, D) using the family assignment method. Dark purple denotes family assignments without the use of a prior on siblingship size, pink denotes family assignments *with* the use of a prior on siblingship size (harmonic mean of siblingship sizes = 1).

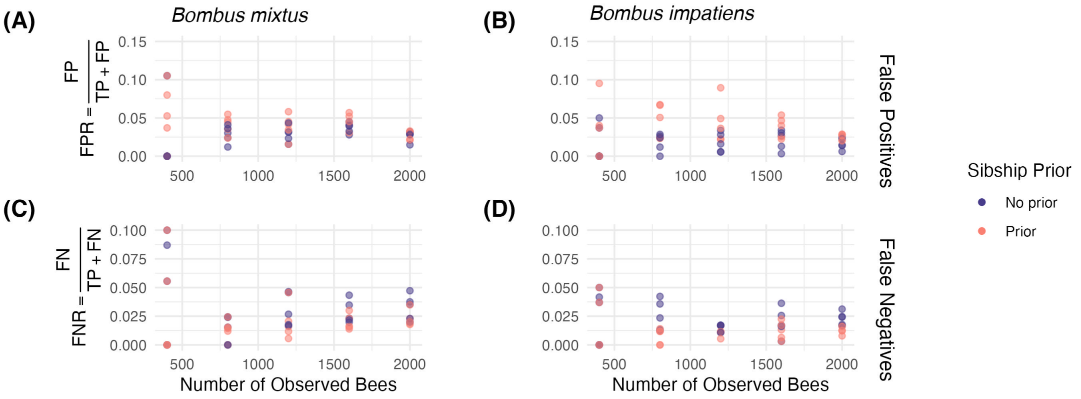


Figure A15: False positive (A-B) and false negative (C-D) rates of siblingship assignment in *B. mixtus* (A, C) and *B. impatiens* (B, D) using the dyad assignment method. Dark purple denotes family assignments without the use of a prior on siblingship size, pink denotes family assignments *with* the use of a prior on siblingship size (harmonic mean of siblingship sizes = 1).

We did not test for statistical significance, but given the qualitative results we decided to utilize the dyad method with a probability threshold of $P = 0.995$ and no siblingship size prior for our final analysis of real data.

##### A2.4.1 Assessing the use of between-landscape sibling exclusion for reducing false positive rates

We next evaluated the use of a sibling exclusion criteria to determine whether this software specifications improves accuracy of siblingship inference.

Previous studies on *Bombus* have varied in their approaches to the spatial scale at which possible siblingships are permitted. Some studies perform separate runs of the software for populations sampled at different sites/regions (Jha & Kremen, 2013) while others group populations at larger scales, permitting the discovery of long distance foraging or dispersal events between sites (Lepais et al., 2010; Mola et al., 2020; Rao & Strange, 2012). While identifying the maximum foraging or dispersal range for different species or landscape contexts is an important goal, we consider two challenges to this method, related to (i) the improbability of capturing individuals engaged in such long distance events, and (ii) the statistical challenges introduced by a very high number of pairwise comparisons. For a more thorough discussion of both points, we direct the reader to Lepais et al. (2010).

Ultimately, we posit that the rate at which long distance foraging events should be captured in the dataset is much lower than both the false positive and false negative rates of the COLONY software. This is due in part to the quadratically increasing search area over which foragers will be dispersed as distance from their nest increases (see discussion in Osborne et al. (2008)). Secondarily, given our large sample size and the spatial structure of our collections, a very high number of pairwise comparisons will occur between individuals at different landscapes (without exclusion). We therefore hypothesized that allowing for siblingship assignments between all individuals would result in a high percentage of false positive relationships that would severely bias estimates of colony locations and foraging behaviour (which are the primary goals of our research).

To test this hypothesis, and to determine whether total sample size had an effect on the utility of excluding between-landscape siblingships, we simulated five datasets of n = 2000 individuals, and subsetted each data set to contain 20, 40, 60, 80, or 100% of the initial samples. These data were simulated to reflect trapping grids which were arranged in a 3 x 2 grid with traps in adjacent grids at least 6km apart (Fig A10 A-B). The minimum distance between adjacent trapping grids in our real data was 5km (also separated by a large river, expected to limit dispersal), and all other landscapes were at least 7km apart. The mean foraging distance of colonies in our simulation was set to 1km (99% of all visitations within 3.32km). This would allow for colonies located midway between trapping grids to sampled at two landscapes, while reflecting the fact that the majority of bumblebee foraging is thought to occur within a few kilometers of the nest. We therefore believe that the simulated data would represent a relatively optimistic view of the number of between-landscape siblingships which could be present in the real data.

We created siblingship exclusion tables for COLONY by excluding (for each individual) all potential siblingships with individuals captured at different trapping grids (landscapes). We ran COLONY on each dataset with and without incorporation of the siblingship exclusion table. Further software specifications for these runs can be found in Table A10. We used the dyad method and an exclusion threshold of $P = 0.995$ as described in the previous section.

We found that for both species (*B. mixtus* and *B. impatiens*) and for all tested sample sizes (n = 400-2000 individuals), between-landscape sibling exclusion resulted in a lower false positive rate (Fig A16 A-B). The variation in false positive rates between independent simulations decreased with increasing sample size. When using between-landscape exclusion, mean false positive rates were fairly consistent across sample sizes, but without between-site exclusion, the mean false positive rate tended to decrease with increasing sample size.

False negative rates were similar for both methods, indicating that between-landscape exclusion did not cause us to lose a high proportion of real between-landscape siblingships (Fig A16 C-D). Indeed, manual inspection of datasets revealed that between-landscape sampling was extremely rare or non-existent, given our parameterization and sample size.

Based on these results, we elected to incorporate a between-landscape sibling exclusion criteria (e.g., only “look" for siblingships between individuals captured on the same trapping grid) in all further analyses. This includes the section above (e.g., Fig A11, A12, A14, A15).

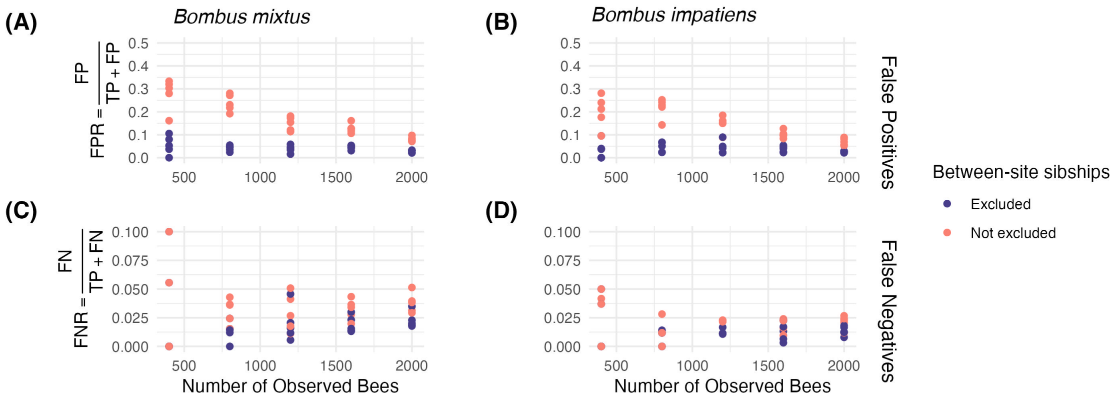


Figure A16: Comparison of between-landscape siblingship exclusion (dark purple) or no exclusion (pink) for minimizing false positive (A-B) and false negative (C-D) rates for *B. mixtus* (A,C) and *B. impatiens* (B,D). Colony assignments were made using the dyad method, with probability threshold = 0.995.

#### A2.4.2 Evaluating the effects of multiple paternity on siblingship inference

Multiple studies have reported low lates of polyandry (queen polygamy) in the North American subgenus *Pyrobombus* (Owen & Whidden, 2013; Payne et al., 2003), which contains both *B. mixtus* and *B. impatiens*. We therefore tested whether our loci are sufficiently informative to identify maternal siblingships (e.g., siblings sharing a mother, but different fathers). Maternal siblings would be colony mates, making them ecologically/behaviourally similar to full siblings, unlike paternal siblings, which would originate from distinct colonies in potentially very different locations. Unfortunately, maternal siblings share on average only 25% of their genomes, in contrast to paternal siblings (50%) and full siblings (75%), making identification of maternal siblingships more challenging that identification of full siblingships.

To test our ability to do so, we simulated datasets of n = 1000 individuals and assigned genotypes under varying assumptions about polyandry rates (0%, 20%, 40%, 60%, 80% and 100% of colonies experience polyandry). We then performed runs of COLONY 2.0.6.5 under both female monogamy and polygamy, to test how these assumptions influence inference. For these analyses, we did not introduce errors or missingness to our simulated genotypes. All colony assignments were made with the family method, as the introduction of half-siblings make the full sibling dyad approach intractable.

When we enforce queen monogamy in COLONY, an increase in the polyandry rate has little or no effect on false positive rates in either species (Fig A17 A-B); increasing the polyandry rate does lead to a steady increase in false negative rates, up to nearly 50% for both species (Fig A17 C-D). This is unsurprising given that enfrocing monogamy prevents us from identifying maternal siblingships. Unfortunately, allowing for queen polygamy in our modelling process does little to alleviate these false negatives (Fig A17 C-D), but results in a substantial increase in the number of false positives (Fig A17 A-B). Around 50-80% of all inferred siblingships are erroneous when we allow for queen polygamy, suggesting that we lack the inferential power to accurately distinguish these relationships.

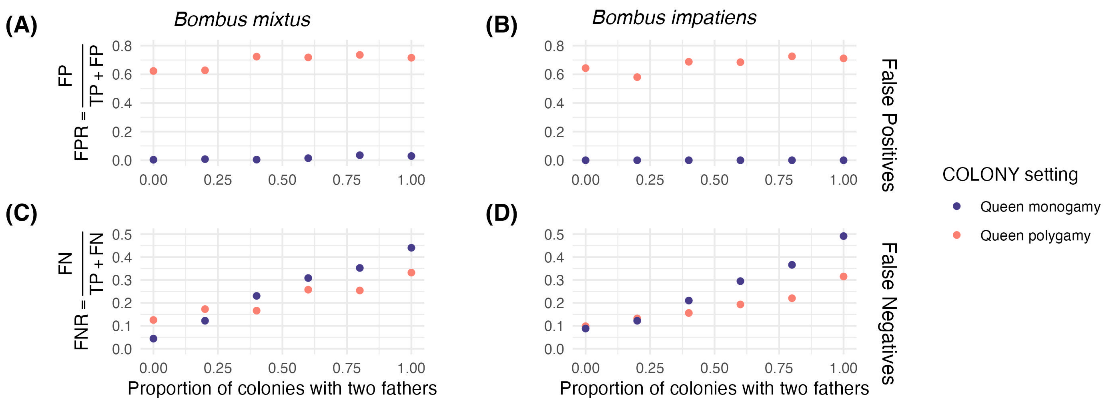


Figure A17: False positive (A-B) and false negative (C-D) from COLONY runs assuming queen monogamy (dark purple) and queen polygamy (pink). A total of 6 datasets were generated for each species (*B. mixtus* (A, C) and *B. impatiens* (C, D)); colony assignments were made twice for each dataset (once under monogamy and once under polygamy).

To further explore this failure, we asked whether a more informative genetic dataset would have allowed us to distinguish maternal siblings. We simulated datasets under varying degrees of polyandry (as above), but this time with a dataset containing 22 loci (e.g., using real allele frequencies from the combined locus sets for *B. mixtus* and *B. impatiens*). With this augmented dataset, we repeated the analyses above and found that while false positive rates were lower for the augmented dataset (Fig A18 A) they were still far above the acceptable range. False negative rates were kept to 10% or less when assuming female polygamy (Fig A18 B), suggesting that while a more informative dataset could help us to identify true maternal siblings, this information would still come at the cost of a very high false positive rate. Further exploration of the accuracy of larger datasets (e.g., SNPs, (Mola et al., 2020)) for determining maternal siblingships is warranted.

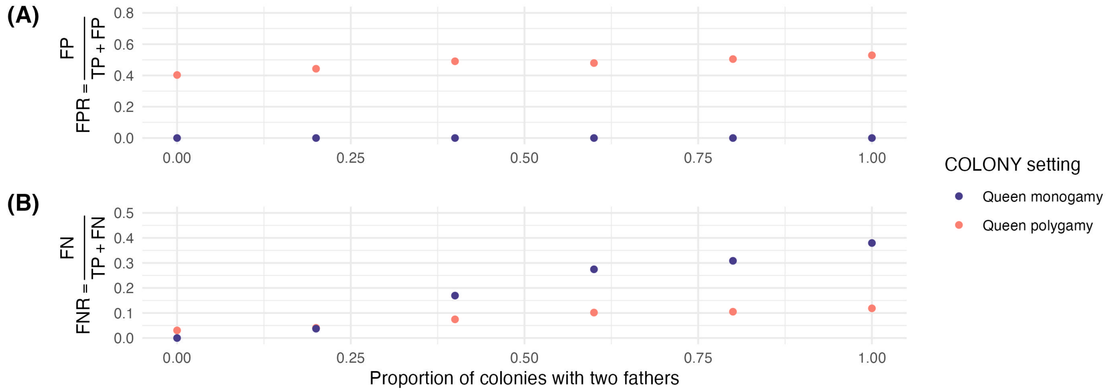


Figure A18: False positive (A-B) and false negative (C-D) from COLONY runs assuming queen monogamy (dark purple) and queen polygamy (pink). A total of 6 datasets were generated for each species (*B. mixtus* (A, C) and *B. impatiens* (C, D)); colony assignments were made twice for each dataset (once under monogamy and once under polygamy).

Table A10: Description of software settings (COLONY 2.0.6.5 (Jones & Wang, 2010)) for simulations.

| **Simulation** | **Comparison** | **Sample Size** | **Sibship Size Prior** | **Between-landscape Exclusion** | **Runs** |
|---|---|---|---|---|---|
| **section 2.2** | Number of COLONY runs | 1200 | yes | yes | 1-5 |
| **section 2.2** | Probability threshold | 1200 | yes | yes | 1-5 |
| **section 2.4** | Siblingship size prior | 400-2000 | yes/no | yes/no | 1 |
| **section 2.2, 2.4** | Families vs dyads | 400-2000 | yes/no | yes | 1-5 |
| **section 2.5** | Between-landscape exclusion | 400-2000 | yes/no | yes/no | 1 |
| **section 2.6** | Mating system | 1000 | no | no | 1 |
| **section 2.6** | Mating system (augmented data) | 1000 | yes | no | 1 |

### A3 Modelling Foraging Distance

To compare foraging distance of the two species, we fitted spatially explicit genetic capture-recapture models following Pope and Jha (2017). These models parametrize the distance decay rate of foraging activity, conditioned on the observation frequency of workers from each colony ($i \in C$) at each transect ($k \in K$). For the purposes of this study, which takes place across multiple landscapes and timepoints, we sum visitation events across sampling rounds and, consistent with our exclusion of siblingships between individuals at different landscapes, we consider visitation events only within the landscape where a colony was observed (e.g., transects $k \in K_l$, where $K_l$ is itself a subset of $K$). The joint likelihood for a simple model where visitation intensity decays exponentially with the square of distance is:

$$\mathbf{y}_i \sim \text{Multinomial}\big(n_i, \mathbf{p}_i\big)$$

$$p_{ik} = \frac{\lambda_{ik}}{\sum_{j=1}^{K_l} \lambda_{ij}}, n_i = \sum_{j=1}^{K_l} y_{ij}$$

$$log(\lambda_{ik}) = -\left(\frac{\|x_k - \delta_i\|}{\rho}\right) + \epsilon_k$$

where $y_{ij}$ is the number of individuals from colony $i$ captured at transect $j$, $\delta_i$ is a nuisance parameter governing the location of colony $i$, $\|x_k - \delta_i\|$ is the Euclidean distance between transect $k$ and colony $i$, $\rho$ is the length scale of foraging activity, and $\epsilon_k$ are transect-year-species intercepts drawn from a common distribution. We incur a sampling offset by adding the log-transformed sampling effort per transect on the linear predictor scale for each $\lambda_{ik}$.

We specify priors $\rho \sim lognormal(log(0.5), 0.5)$, $\epsilon \sim \mathbb{N}(0, \sigma^2)$, and $\sigma \sim exponential(1)$. In our initial models, $\delta$ received an implicit uniform prior as in Pope and Jha (2017); to speed up computation, we limited the bounds on this prior to ±5km easting/northing from the trapping grid where the colony was observed. We conducted model inference using Stan (Carpenter et al., 2017).

Pope and Jha (2018) propose that capture records of siblings along transects contain more information regarding *relative* foraging distance (e.g., comparisons between landscapes or species) than *absolute* foraging distance, likely due to the low recapture rate of each colony. Indeed, our initial estimates of absolute foraging length scale, $\rho$, were sensitive to priors on colony locations $\delta$.

To better understand this model pathology, we conditioned the model on three datasets: (1) an "ideal" simulated dataset, consisting of 562 colonies with an average siblingship size of 17.8 individuals (Fig A19A), (2) a biologically realistic simulated dataset, consisting of 1390 colonies with an average siblingship size of 1.4 (Fig A19B), and (3) data collected in the field for *B. impatiens*, consisting of 2632 colonies with average siblingship size of 1.3 (Fig A19C). Both simulated datasets were produced with a data-generating value of $\rho = 0.3$ km (e.g., 99% of foraging occurs within approx. 1380m of the nest).

The inferred posterior means for $\rho$ were 0.296 (CI: $[0.289, 0.302]$) (ideal simulated dataset), 0.468 (CI: $[0.426, 0.515]$) (biologically realistic simulated data), and 0.354 (CI: $[0.323, 0.388]$) (*B. impatiens* dataset) (Fig A19 D-F). While the model conditioned on the ideal dataset is able to pinpoint the true data generating value, a more biologically realistic sibling recapture appears to result in overestimation of the foraging range. This overestimation is systematic (data not shown, but see Pope and Jha (2017)).

We next produced pairs plots of the posterior samples for $\rho$, $\sigma$, and the unnormalized log joint posterior density (not shown) to explore the relationship between these parameters. For models conditioned on biologically realistic simulations and real datasets, $\rho$ was negatively correlated with the unnormalized log joint posterior density. To more clearly visualize this pattern, we fixed $\rho$ across a range of values and computed the optimum of the log joint posterior density (similar to a profile likelihood, but incorporating priors) (Fig. A19 G-I). We observed that the joint density was largest around the data-generating value of $\rho$, and decreased at larger and smaller values. When, then, did our full Bayesian model systematically overestimates $\rho$? Recall that the support for a model is the product of the likelihood and the area over which the likelihood is integrated — while the joint density may be maximized for small $\rho$, small values of $\rho$ are restrictive on other terms in the model (e.g., the high-dimensional vector of latent colony locations, $\delta$). This reduces the volume of the parameter space which is consistent with the data. Larger $\rho$ values can achieve greater posterior mass because their lower likelihood is compensated by greater volume in the latent space. This phenomenon is analagous to the typical set in high dimensions, as described in Betancourt (2018). In this case, an ideal dataset provides enough information to penalize large $\rho$ values and result in accurate posterior inference, but for less informative (realistic) datasets, an informative prior on $\delta$ is necessary to constrain the volume of the parameter space consistent with the data and achieve accurate estimation of $\rho$.

To achieve this, we apply a bivariate Gaussian prior on $\delta_i$ which is centered at the middle of the replicate landscape where colony $i$ is observed.

$$\delta = \begin{pmatrix} \delta_x \\ \delta_y \end{pmatrix} \sim \mathcal{N}\Big(\mathbf{0}, \sigma_d \mathbb{I}\Big)$$

This prior should not be interpreted biologically (e.g., “bumble bees are more likely to nest at the center of our study landscapes because they have an affinity for pesky graduate students in high-vis vests") but rather as a prior on the observational process. The colonies we detect are only a subset of all colonies located on the landscape — indeed, if we fit the distribution of siblingships sizes (Fig A19 C) to a truncated Poisson to estimate the zero-class, we find that the majority of colonies are not observed at all. While colonies in general may be uniformly distributed, we are most likely to observe those near the center of the trapping grid, as we have surveyed a larger proportion of their foraging kernel than we have for colonies located on the periphery.

To test the estimability of relative foraging distance, we fitted our updated model to data simulated with a range of $\rho$ values and performed sensitivity analysis using biologically informed priors $\sigma_d \in (0.5, 1, 1.5)$ km. For a bivariate Gaussian, the distribution of distances from the mean (center of the replicate landscape) is a Rayleigh distribution with scale parameter $\sigma = \sigma_d\sqrt{2 - \pi/2}$. This prior places 99% of probability within $3\sigma_d$ km of the landscape center. Using this prior, we observed differences in $\rho_I$ (model-estimated $\rho$) over a range of biologically realistic $\rho_G$ (data-generating $\rho$) (Fig A20), confirming the utility of biologically realistic siblingship capture-recapture datsets for estimating relative foraging distance. Unfortunately, even with a more informative prior on $\delta_i$, we observed that inference of absolute foraging distance is sensitive to the specification of this prior (weaker priors tend to produce larger estimates of $\rho$).

The foraging distance model we present in the main body of the text utilizes $\sigma_d = 1$ km, which places 99% of the prior probability for colony locations within 3 km of the landscape center. This is a reasonable choice given that our trapping grids are roughly 1.7 x 1.7 km, meaning that latent colony locations colonies should be constrained to fall within approx. 2km of all transects (with a higher *a priori* probability of being located closer to the center of the grid). We tested the sensitivity of our results to this prior choice, and found that while absolute estimates of $\rho$ were sensitive to $\sigma_d$, relative estimates (e.g., comparisons between species) were relatively consistent (Fig A21). We do note that the 95% credible intervals for the difference in $\rho$ between species marginally overlapped with zero when $\sigma_d = 0.5$ km and $\sigma_d = 1.5$ km (97.0% and 95.9 % of posterior draws for the difference were $\geq 0$, respectively). These two priors equate to colonies falling within 1.5 km and 4.5 km of the landscape center with 99% probability and therefore represent relatively extreme assumptions about the distribution of colony locations. We therefore feel confident in our conclusion that *B. impatiens* has a larger foraging distance than *B. mixtus*, but acknowledge that the absolute estimates of $\rho$ are sensitive to this prior choice and should be interpreted with caution.

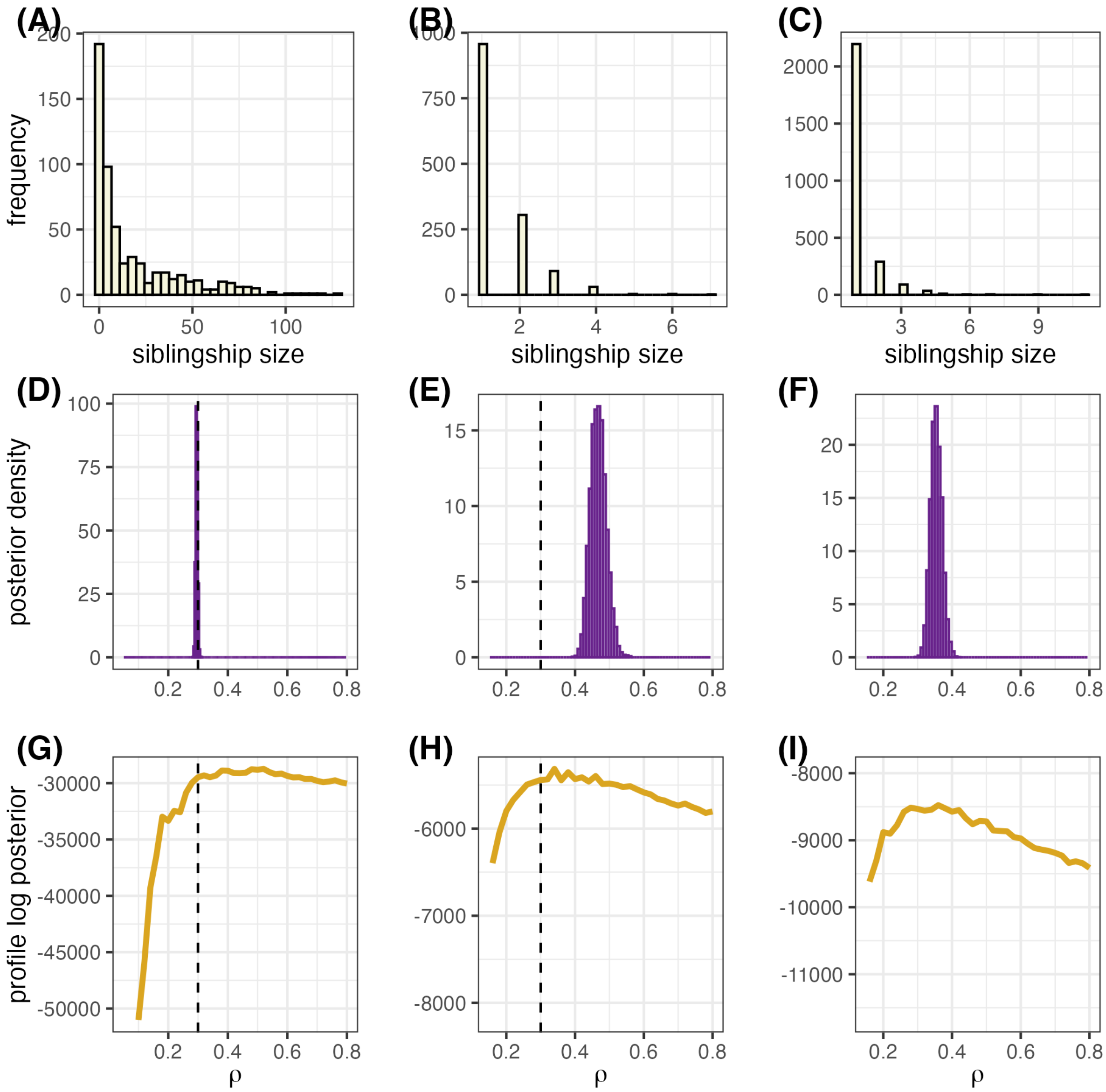


Figure A19: Conditioning a simple foraging distance model on simulated and real bumblebee spatial mark-recapture datasets. We perform inference using the initial model and three datasets: an ideal simulated dataset (A, D, G), with an average siblingship size of 37 individuals per colony, a biologically realistic simulated dataset (B, E, H) with an average siblingship size of 1.4 individuals, and an empirical dataset for *B. impatiens* (C, F, I) with an average siblingship size of 1.3 individuals. Panels A-C show the distribution of siblingship sizes for each dataset; D-F show the posterior distributions of $\rho$, the foraging length scale; G-I show the optimal unnormalized log posterior for fixed values of $\rho$. The vertical dashed lines in D, E, G, and H represent the data-generating $\rho$ used for simulations.

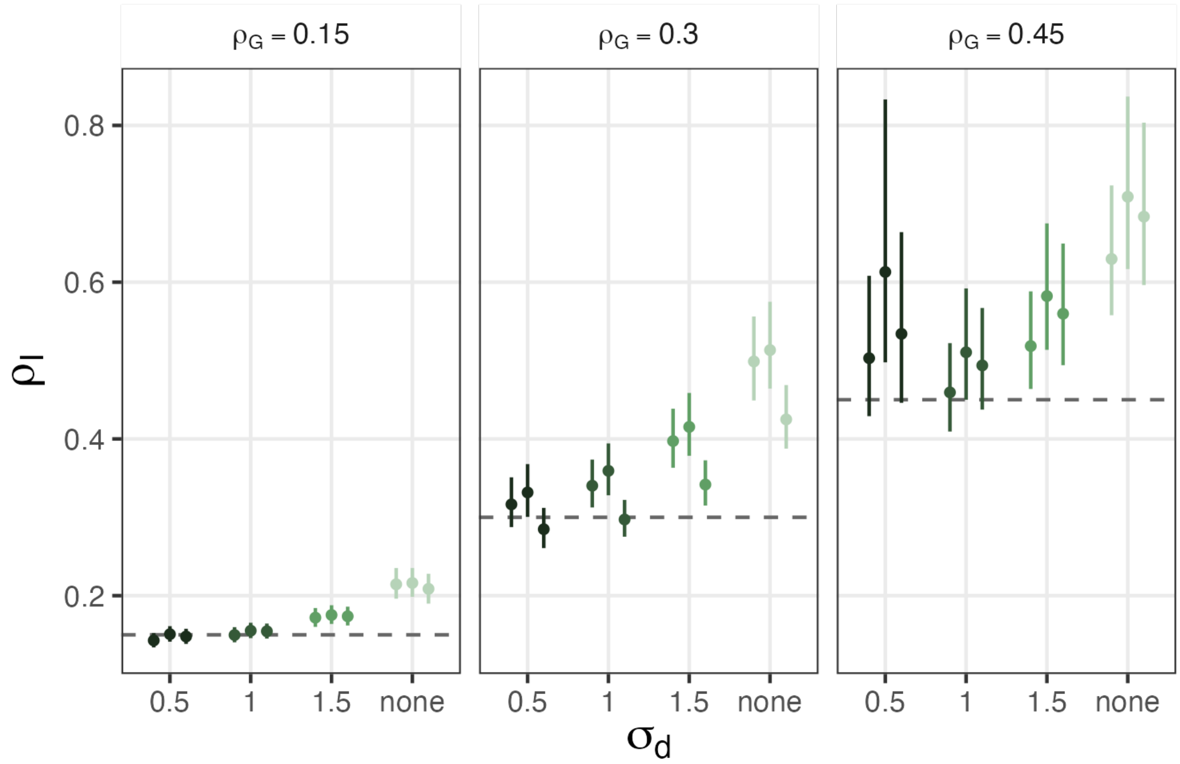


Figure A20: Model-estimated foraging length scale ($\rho_I$) across a range of data-generating foraging length scales ($\rho_G$) and priors on colony locations ($\sigma_d$). The dashed line represents the true value of $\rho_G$ for each facet. Models with $\sigma_d$ = "none" were given an implicit uniform prior on colony location, limited to within ±5 km Northing/Easting of the trapping grid where they were observed.

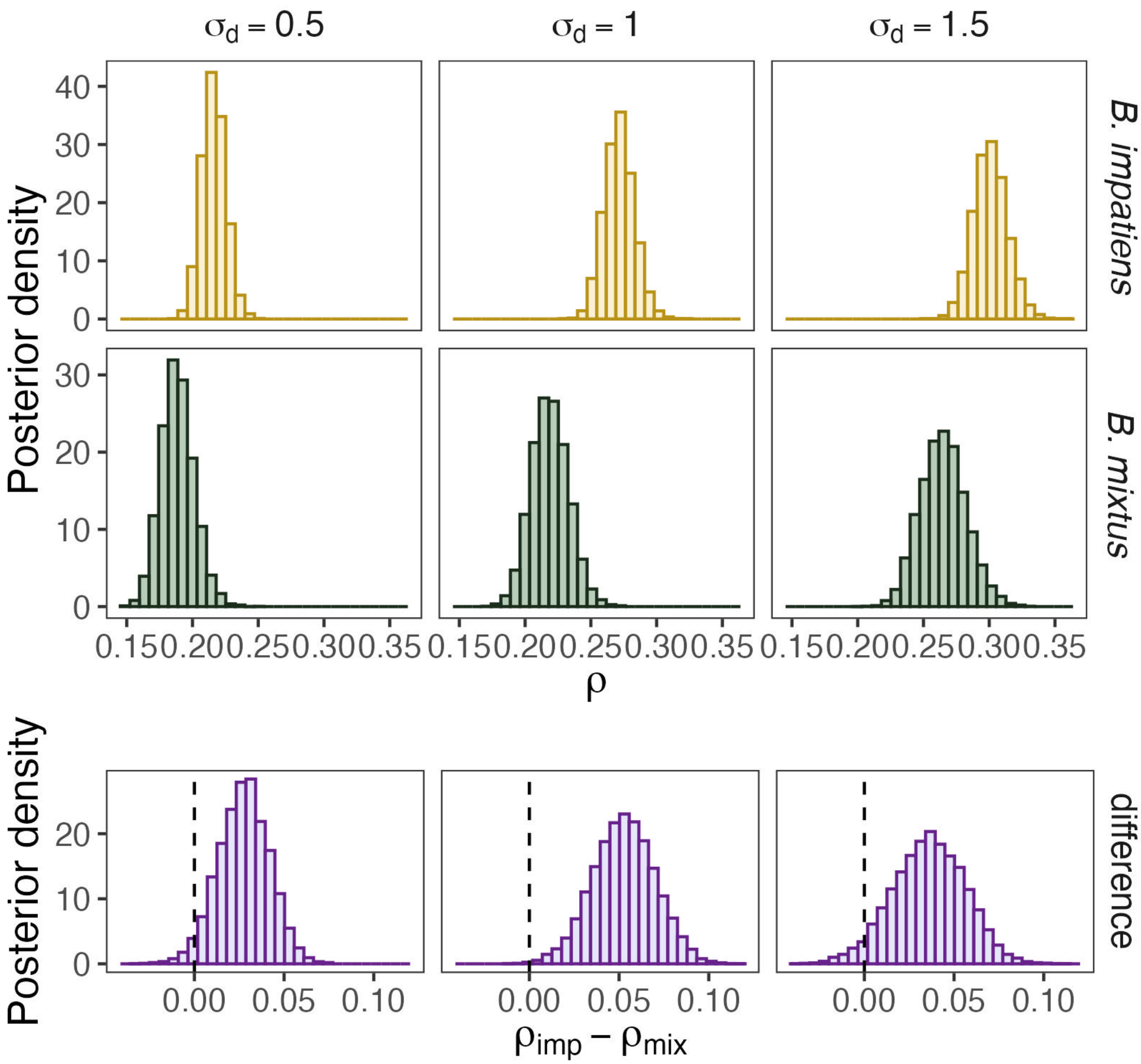


Figure A21: Sensitivity analysis of foraging distance estimates to priors on colony locations. The first two rows of plots show the posterior distribution of $\rho$ for *B. mixtus* (green) and *B. impatiens* (yellow) for three different priors on colony locations ($\sigma_d$ = 0.5, 1, and 1.5 km). The bottom row shows the posterior distribution of the difference in $\rho$ between species for each prior choice. Vertical dashed lines indicate zero.
